# Chronosequence of invasion reveals minimal losses of population genomic diversity, niche expansion, and trait divergence in the polyploid, leafy spurge

**DOI:** 10.1101/2023.04.04.535556

**Authors:** Thomas A. Lake, Ryan D. Briscoe Runquist, Lex E. Flagel, David A. Moeller

**Affiliations:** Department of Plant and Microbial Biology, University of Minnesota, St. Paul, Minnesota, USA; Gencove, Long Island City, New York, USA

**Author notes:** Correspondence Thomas A. Lake and David A. Moeller, Department of Plant and Microbial Biology, University of Minnesota, St. Paul, Minnesota, USA. **Emails and ORCID,** Thomas Lake, Ryan Briscoe Runquist, Lex Flagel, David Moeller.

**Keywords:** : adaptation, colonization bottlenecks, plant invasion, population genetic structure, range shift, rapid evolution

## Abstract

Rapid evolution may play an important role in the range expansion of invasive species and modify forecasts of invasion, which are the backbone of land management strategies. However, losses of genetic variation associated with colonization bottlenecks may constrain trait and niche divergence at leading range edges, thereby impacting management decisions that anticipate future range expansion. The spatial and temporal scales over which adaptation contributes to invasion dynamics remains unresolved. We leveraged detailed records of the ∼130 year invasion history of the invasive polyploid plant, leafy spurge (*Euphorbia virgata*), across ∼500km in Minnesota, U.S.A. We examined the consequences of range expansion for population genomic diversity, niche breadth, and the evolution of germination behavior. Using genotyping-by-sequencing, we found some population structure in the range core, where introduction occurred, but panmixia among all other populations. Range expansion was accompanied by only modest losses in sequence diversity, with small, isolated populations at the leading edge harboring similar levels of diversity to those in the range core. The climatic niche expanded during most of range expansion, and the niche of the range core was largely non-overlapping with the invasion front. Ecological niche models indicated that mean temperature of the warmest quarter was the strongest determinant of habitat suitability and that populations at the leading edge had the lowest habitat suitability. Guided by these findings, we tested for rapid evolution in germination behavior over the time course of range expansion using a common garden experiment and temperature manipulations. Germination behavior diverged from early to late phases of the invasion, with populations from later phases having higher dormancy at lower temperatures. Our results suggest that trait evolution may have contributed to niche expansion during invasion and that distribution models, which inform future management planning, may underestimate invasion potential without accounting for evolution.

## Introduction

Invasive species experience considerable changes to genetic variation during the process of introduction and subsequent invasion (Lee 2002; Suarez and Tsutsui 2008). In particular, it has been well-documented that founder effects and genetic drift can cause substantial losses of genetic diversity during the colonization process (Dlugosch and Parker 2008; Uller and Leimu 2011). Following initial establishment, further losses of variation may occur during range expansion. However, the magnitude of changes in genetic variation depends upon the number of introductions and the severity of population bottlenecks (Nei et al. 1975; Uller and Leimu 2011; Welles and Dlugosch 2019). Such changes in genetic variation early in the invasion process may influence the capacity for adaptation, forecasts of range expansion, and subsequent management decisions.

Following colonization, some non-native species exhibit rapid population growth and dispersal into new environments (Sakai et al. 2001). The process of invasion is often highly variable, involving repeated founder events and density-dependent population growth, especially Allee effects (Melbourne and Hastings 2009; Sullivan et al. 2017). In addition to affecting the speed of invasion, these population fluctuations can influence levels of genetic diversity and structure across an invaded range (Austerlitz et al. 1997; Excoffier 2004). For example, range expansion is expected to cause a reduction in allelic richness and heterozygosity with increasing distance from the origin of expansion (Slatkin and Excoffier 2012; Peter and Slatkin 2013). The prevalence of drift during invasion may also cause populations to depart from migration-drift equilibrium, resulting in a lack of isolation-by-distance (Wright 1943; Slatkin 1987, 1993; Hutchison and Templeton 1999). Last, mutations arising at the range edge may rise in frequency due to genetic drift, “surf” along the expanding front, and travel long distances (Klopfstein et al. 2006; Excoffier and Ray 2008). Models of allele surfing indicate that rapid range expansion can produce clinal variation in allele frequencies and increase the frequency of loci with private alleles (Klopfstein et al. 2006; Excoffier and Ray 2008; Goodsman et al. 2014). Such clines can emerge for any type of mutation (beneficial, neutral, deleterious), and therefore could reflect drift and/or selection (Lehe et al. 2012; Peischl et al. 2013; Koski et al. 2019). Overall, the prevalence of drift during range expansion has the potential to influence the capacity for adaptation as organisms encounter novel environments, particularly when functionally-important allelic variation is lost.

An increasing body of evidence suggests that rapid phenotypic evolution can be important to the process of range expansion (Colautti and Barrett 2013; Hodgins et al. 2018; Selechnik et al. 2019; Ma et al. 2020). Forecasts of range expansion in invasive species can underpredict the potential extent of invasion if they assume a species does not evolve or adapt over short time scales (Chardon et al. 2020; Collart et al. 2020). A recent meta-analysis indicated that the signature of local adaptation in invasive species was at least as strong as in native species, even when accounting for variation in life history (Oduor et al. 2016). Invasive species frequently expand across strong environmental gradients and into novel niche space (Atwater et al. 2018; Bates and Bertelsmeier 2021). Such niche expansion may require adaptive evolution at the invasion front (Chown et al. 2015; Moran et al. 2017; Hodgins et al. 2018). For example, purple loosestrife rapidly diverged in flowering time and plant size during invasion across latitudinal gradients in growing season length (Colautti and Barrett 2013). Despite evidence of rapid evolution in some systems, range expansion may not involve any changes in the organism’s niche if the only limit to spatial expansion is dispersal and time. As such, the apparent expansion of the climate niche with invasion may not actually involve the evolution of ecologically-important traits. While tests of local adaptation within an invaded range remain few, there is growing appreciation that rapid evolution is likely to shape the trajectory of range expansion in non-native species (Hodgins et al. 2018; Woods and Sultan 2022).

Phenotypic evolution during range expansion may be caused by spatially-variable selection and/or neutral processes (e.g., spatial sorting) (Keller and Taylor 2008; Colautti and Lau 2015; Hodgins et al. 2018). Adaptive evolution, in particular, may be paramount to the invasion process if selection in response to novel environments results in trait changes that enhance a species’ capacity to establish in new habitats (Prentis et al. 2008; Williams et al. 2016; Hodgins et al. 2018; Woods and Sultan 2022). While reciprocal transplant experiments are the gold standard for testing for adaptation, the translocation of invasive species for these experiments is subject to ethical concerns and legal restrictions in many areas. Alternatively, researchers have started to use ecological niche models (ENMs) to identify important environmental gradients that span from optimal to marginal habitat, such as from a range core to edge. Predictions are then made about traits that may promote adaptation to the novel environments found in marginal habitats (Searcy and Shaffer 2016; Dixon and Busch 2017; Capblancq et al. 2020; Morente-López et al. 2022). Finally, common garden experiments can determine whether the putative traits under selection have differentiated across the key environmental gradients identified by ENMs. Taken together, this series of approaches can provide insight into the role of adaptation in the process of niche expansion at leading range edges.

Among invasive plant species, polyploidy is prevalent (Pandit et al. 2011) and can influence the process of range expansion (Van de Peer et al. 2021). The frequency of polyploid species increases with higher latitudes, lower temperatures, and seasonally drier environments (Brochmann et al. 2004; Rice et al. 2019). Direct effects of polyploidy on physiological, morphological, and phenological traits may facilitate niche shifts (Glennon et al. 2014; Blaine Marchant et al. 2016; Brittingham et al. 2018; Wang et al. 2022) and preadapt polyploids to new environments (Treier et al. 2009; Lachmuth et al. 2010). Polyploidy may also influence the capacity for adaptation during range expansion as new environments are encountered (te Beest et al. 2012; Baniaga et al. 2020). Although genetic drift during range expansion can cause losses of genetic diversity, drift may have less severe effects (e.g., inbreeding depression) in polyploids relative to diploids when there is polysomic inheritance (Moody et al. 1993; Soltis and Soltis 2000). Despite numerous polyploid invaders, they have been the subject of few studies because of substantial challenges with the application of evolutionary genetic analyses that were developed for diploids (Rutland et al. 2021).

In this study, we used a well-documented chronosequence of invasion to examine the consequences of range expansion for population genomic diversity, climatic niche breadth, and the evolution of germination behavior in the polyploid, leafy spurge (*Euphorbia virgata*). We focused on one area of introduction to southwestern Minnesota, U.S.A and subsequent range expansion to the north and east. We were interested in examining the severity of losses of genetic diversity following introduction and its potential consequences for invasion, particularly since existing species distribution models predict a low probability of range expansion at the current leading range edge (Lake et al. 2020). Introduction to this region occurred in the 1890s in southwestern Minnesota with subsequent range expansion to northeastern Minnesota (ca., 500 km), where populations are currently rare, isolated, and small. Based on a detailed historical occurrence dataset, we defined a range core, area of early expansion, area of late expansion, and invasion front.

First, we sampled populations along the gradient from core to invasion front using two sampling schemes to quantify population genomic diversity and structure (using reduced representation sequencing). Samples of multiple individuals from 14 populations (population samples) allowed us to quantify changes in sequence diversity among populations over the course of range expansion. Samples of single individuals from 157 populations (landscape samples) allowed us to test for fine-scale population structure (e.g., isolation-by-distance) over the time series of range expansion. Second, we tested whether range expansion involved niche expansion - i.e., occurred into novel climatic environments. Third, we developed an ecological niche model (ENM) to test whether habitat suitability declines from range core to invasion front and to determine which environmental gradients are most strongly associated with high versus low habitat suitability. Warm season temperature had the greatest positive contribution to habitat suitability. Because temperature is known to modulate germination behavior in leafy spurge, and because establishment in new habitats is dependent upon successful germination timing, we focused on this trait for common garden experiments. Past work has also suggested that shifts in seed dormancy might facilitate invasion at leading range edges (Mathias and Kisdi 2002; Travis et al. 2021). We examined the responses of seeds from early versus late in invasion to five temperature regimes in a common garden experiment. We specifically tested whether there was an interaction between geographic region (early vs. late invasion stages) and temperature regime, which would indicate divergence in germination behavior over the course of invasion.

## Methods

### Invasion and natural history of leafy spurge

#### Natural history

Leafy spurge has invaded approximately two million hectares across the northern tier of the United States and southern Canada (Duncan et al. 2004). While it is most commonly found in dry, open sites with well-drained soils (e.g., prairies), it can occasionally occur in seasonally wet meadows and riparian areas (Selleck et al. 1962). Leafy spurge impacts rangelands and natural habitats by competitively displacing native species (Hein and Miller 1992). When damaged, plants exude a toxic white latex that deters grazing (Lym and Kirby 1987; Lym 1998).

Leafy spurge spreads locally via rhizomes and ballistic seed dispersal (Morrow 1979). Longer distance dispersal has been proposed to occur via animals or agricultural machinery (Pemberton 1988; Lacey et al. 1992). Seeds germinate in spring or may remain dormant in soil for at least two years (Hanson and Rudd 1933; Selleck et al. 1962). Flowers are insect pollinated and the mating system is primarily outcrossing (Selleck et al. 1962).

Leafy spurge is an auto-allohexaploid that likely originated from hybridization between closely-related *Euphorbia* species, although the progenitor species are not yet known (Schulz-Schaeffer and Gerhardt 1989; Riina et al. 2013). It has been the subject of several genomic investigations but lacks a full genome assembly and annotation (Chao et al. 2005; Horvath et al. 2015, 2018; West et al. 2023).

#### Invasion history

One introduction of leafy spurge occurred into southwestern Minnesota ca., 1890 purportedly via contaminated grains imported from southern Russia (Batho 1932; Hanson and Rudd 1933; Dunn 1985). Following introduction, the range expanded to eastern South Dakota by ca., 1902 (Bakke 1936), eastern North Dakota by ca. 1909 (Hanson and Rudd 1933), and southern Manitoba and Saskatchewan by the 1920s (Batho 1932; Selleck et al., 1962). Hanson and Rudd (1933) documented in detail the distribution of leafy spurge across Minnesota and neighboring regions, providing a baseline for understanding the timeline of subsequent range expansion (Figure S1). By the late 1970s, leafy spurge had become common throughout grasslands of the north-central plains (Dunn 1979, 1985). Invasion of the boreal forest region of northeastern Minnesota began in the 1940s and 1950s with isolated occurrences (Lakela 1965) and populations were not common until the 1990s. This invasion front has persisted with limited expansion further northeast.

#### Delineating the timeline of range expansion

We digitized the earliest known point record map (Hanson and Rudd 1933) using ArcGIS Pro (ESRI, 2022). We then applied empirical Bayesian kriging to produce a continuous density surface that represented the density of populations in the north-central plains. From this density surface, we applied an equal-interval threshold to demarcate a range core, area of early expansion, area of late expansion, and invasion front that corresponded to four density categories across Minnesota and surrounding states (Figure 1A). We verified these demarcations with published accounts of the invasion history (described above).

**Figure 1.**
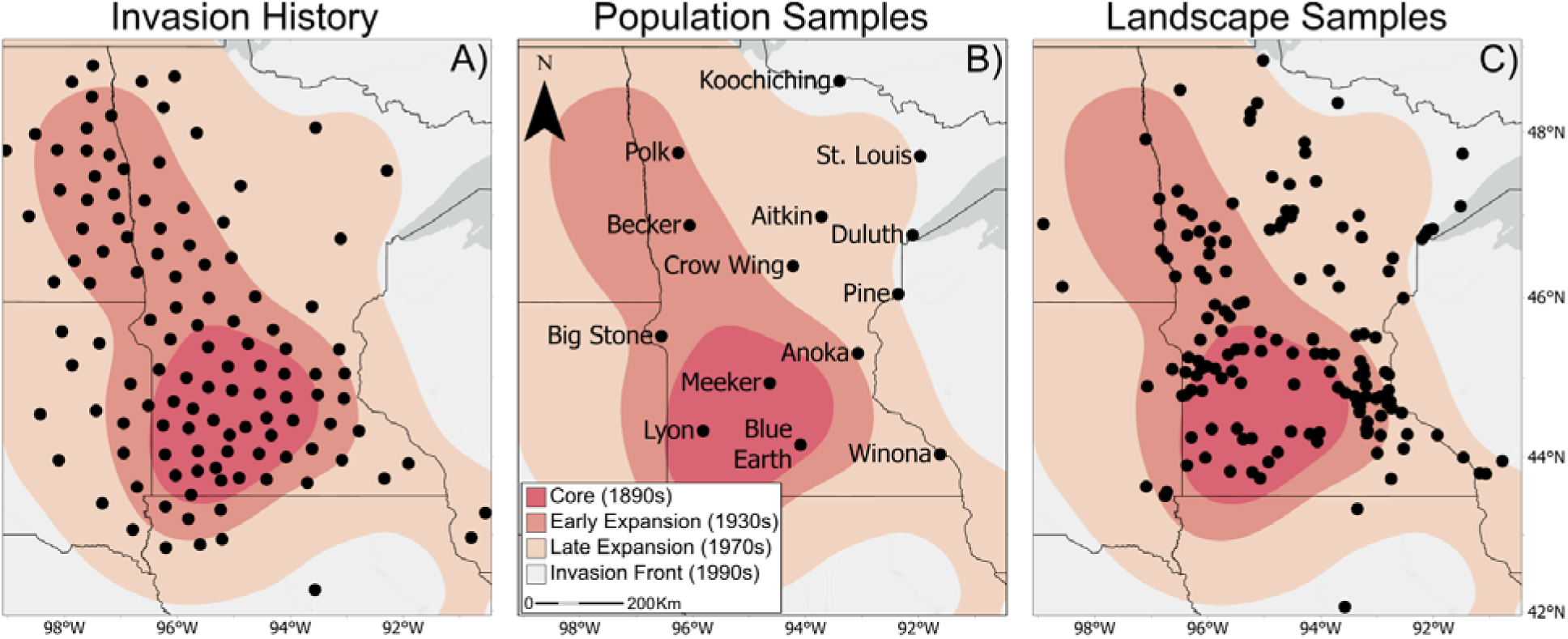
A) Invasion history reproduced from Hanson & Rudd (1933). The map was used to delineate four phases of invasion: core, early expansion, late expansion, and invasion front. Tissue collection sites for B) 14 population samples and C) 157 landscape samples of leafy spurge.

### Population genomic diversity and structure

#### Sampling and sequencing

In 2019, we collected leaf tissue from six individuals in each of 14 populations distributed evenly across Minnesota (*hereafter:* population samples; Figure 1B). In addition, we collected tissue from one individual in each of 157 populations distributed relatively evenly across Minnesota, eastern South Dakota, eastern North Dakota, and western Wisconsin (*hereafter:* landscape samples; Figure 1C). We sampled tissue from individuals that were at least five meters apart to minimize collecting from the same genet and placed tissues immediately in silica for preservation until DNA extraction (Table S1).

We extracted DNA using QIAGEN DNeasy Plant Mini Kits (QIAGEN Inc.). Dual-indexed GBS (genotyping-by-sequencing) libraries were created using the BamHI + NsiI enzyme combination. All libraries were pooled and sequenced on an Illumina NovaSeq System (Illumina Inc., San Diego, CA, USA) with 1x100-bp sequencing. Once sequenced, the reads were demultiplexed and balanced with a mean quality score ≥ Q30 for all libraries. We filtered low-quality bases using Trimmomatic (Bolger et al. 2014) and used Stacks v.2.5.9 (Rochette et al. 2019) to build loci *de novo* (i.e., without aligning reads to a reference genome). Overall, we obtained 510 million reads across the 241 samples (599,386 – 3,376,078 of raw reads per individual). Mean read depth per locus ranged from 14x to 26x (Supplementary Methods).

We called SNPs using polyRAD v.2.0.0 (Clark et al. 2019), a Bayesian algorithm designed for polyploid GBS data. PolyRAD estimates the genotype probabilities for each individual from read depth distributions with ploidy level as a prior. First, we filtered our dataset using the H_ind_/H_e_ statistic to cull likely paralogous loci (Clark et al. 2022). Next, we estimated posterior probabilities for each genotype using the ‘IterateHWE’ function with Hardy-Weinberg equilibrium as the prior (Gerard and Ferrão 2020; Clark et al. 2022). For each individual at each locus, we exported the most probable genotype for subsequent analyses (Supplementary Methods).

We implemented a second filtering step to account for potential biases caused by homoeologous loci present in our dataset. Because leafy spurge is an auto-allohexaploid (Schulz-Schaeffer and Gerhardt 1989), we expect homoeologous loci to have a 2:1 allelic ratio (e.g. AAAABB genotype; Horvath et al. 2018). Indeed, we identified a peak in the minor allele frequency spectrum around 0.33 (Figure S2). We removed loci with a minor allele frequency above 0.26 from our dataset because they are likely to have an excess of homoeologous genotypes (Figure S2). While essential, this second filtering step limits our capacity to understand absolute levels of sequence diversity. However, our primary goal was to examine changes in sequence diversity over the course of invasion rather than absolute quantities. After filtering, 3,176 loci remained for downstream analyses.

#### Population structure

We performed an analysis of molecular variance (AMOVA) to quantify the proportion of genetic variation partitioned among populations, among individuals within populations, and within individuals (Excoffier et al. 1992; Meirmans 2012, 2020). We estimated genetic variance components using the rho statistic (Ronfort et al. 1998) and determined significance using permutation tests (n= 999) using the R package poppr v.2.8.6 (Kamvar et al. 2015). Further, we checked for clonality among samples using the ‘clonecorrect’ function in (Kamvar et al. 2015).

We assessed population structure using the Bayesian clustering algorithm STRUCTURE v.2.3.4 (Pritchard et al. 2000). We ran the analysis 10x for each of K = 1-10 with 500,000 Markov Chain Monte Carlo iterations with a 50,000-run burn-in period, specifying an admixture model with the assumption of uncorrelated allele frequencies. We used ‘structure_threader’ in Python 3 (Pina-Martins et al. 2017) to parallelize runs across clusters (K = 1 – 10). We determined the most plausible number of clusters using the ΔK method (Evanno et al. 2005) and STRUCTURE HARVESTER web v.0.6.94 (Earl and vonHoldt 2012).

We performed principal component analyses (PCA) to examine population structure using GENODIVE v.3.0.4 (Meirmans 2020). We performed PCA separately for the population samples and the landscape samples.

We tested for isolation by distance (IBD; Wright 1943) in the population samples by estimating genetic differentiation as G_ST_ (Dufresne et al. 2014) using GENODIVE v.3.0.4 (Meirmans 2020). We also tested for IBD in the landscape samples by calculating Nei’s genetic distance (D; Nei 1972) (Meirmans 2020). We subset the landscape samples according to successive stages of range expansion and tested for IBD within each subset (i.e., within range core, then successively including areas of early expansion, late expansion, and invasion front). We tested for a relationship between the pairwise genetic and geographic distance (kilometers) matrices using Mantel tests with 9,999 permutations in the R package ‘adegenet’ v.2.1.8 (Jombart 2008).

#### Tests for changes in population genetic diversity during range expansion

For each of the 14 population samples, we estimated the inbreeding coefficient (G_IS_) and observed heterozygosity (H_o_; gametic heterozygosity; Moody et al. 1993), which corrects for potential overestimates of heterozygosity in polyploids by calculating the fraction of heterozygotic gametes for each genotype, using GENODIVE v.3.0.4 (Meirmans 2020). We also calculated the number of private alleles (P) using poppr v.2.8.6 (Kamvar et al., 2014). We calculated Tajima’s *D* (Tajima 1989) using DNASp v.6.0 (Rozas et al. 2017) to gauge if populations have an excess or deficit of rare alleles, which can be indicative of population expansion following a bottleneck (negative *D*) or sudden population contraction (positive *D*), respectively. Because we filtered loci with higher minor allele frequencies, Tajima’s *D* should be biased to lower values. Finally, we tested for evidence of genetic bottlenecks based on linkage disequilibrium using NeEstimator v.2.0.1 (Do et al. 2014).

We tested for changes in population genetic parameters (H_o_, G_IS_, P, and Tajima’s *D*) as populations dispersed beyond the range core (Table 1). For each statistic separately, we used a multiple linear regression that included latitude, longitude, and their interaction as independent variables with the R package ‘car’ v.3.1-2 (Weisberg, 2019). As all late expansion and invasion front populations are located either north or east of the range core, latitude and longitude describe the northern and eastern invasion spread, respectively. In the model of private alleles, we identified the ‘Winona’ population as an outlier using diagnostic plots of residuals (Figure S3), so we removed it from the analysis. We were unable to conduct such a geographic test of N_e_ from the NeEstimator analyses because all but three populations had an N_e_ of infinity (Table S2).

**Table 1.**
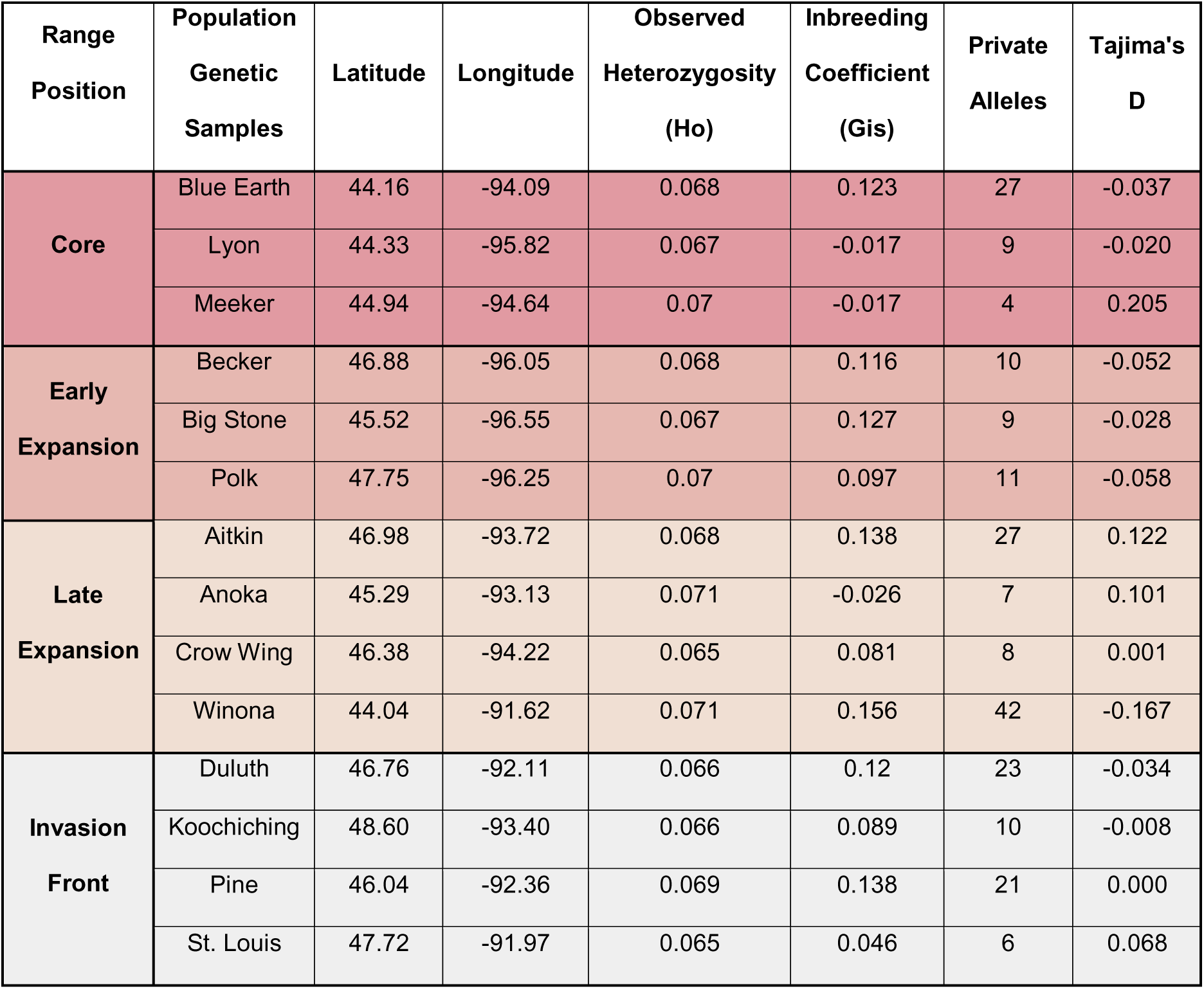
Locality information and genetic diversity metrics (observed heterozygosity, Ho; inbreeding coefficient, G_is_; number of private alleles, and Tajima’s D) for 14 leafy spurge population samples from four phases of range expansion (see Figure 1).

### Niche breadth and habitat suitability

#### Environmental data

We downloaded 19 bioclimatic variables at a 30 arcsecond resolution (∼1 km) from Worldclim (http://worldclim.org/version2). We used a principal component analysis based on the 157 landscape samples to examine patterns of correlation among bioclimatic variables. The biplot of the first two principal components (74.7% variance explained) showed that there were three sets of correlated variables: one set includes mostly precipitation variables, a second set includes mostly temperature variables, and a third set includes variables that describe seasonality and temperature ranges (Figure S4). Because the selection of one variable from a set of correlated ones is challenging (e.g., choosing one precipitation variable from a set of correlated precipitation variables), studies have advocated for choosing variables that are biologically most relevant to the organism and largely uncorrelated (Chapman et al. 2017; Petitpierre et al. 2017).

We selected minimum temperature of the coldest month (Bio 6), mean temperature of the warmest quarter (Bio 10), and precipitation of the warmest quarter (Bio 16). These three variables provide biologically meaningful axes of climate variation that are relevant to key life history stages for an herbaceous perennial (Petitpierre et al. 2017; Chapman et al. 2017). Specifically, minimum cold temperatures are important for overwintering and cold tolerance (Chapman et al. 2017), whereas the temperature of the warmest quarter likely influences seed germination, plant growth, and phenological transitions (Wolkovich et al. 2013). Precipitation of the warmest quarter likely also affects plant growth and reproduction, with lower precipitation associated with reduced growth and increased drought stress in northern temperate ecosystems (Petitpierre et al. 2017; Gorton et al. 2019).

#### Tests for niche differentiation during range expansion

We tested for climatic niche differentiation between the range core, early expansion, late expansion, and invasion front using the ‘ecospat’ R package v.3.5.1 (Di Cola et al. 2017). We sampled the total extent of the background environmental space with a principal component analysis using 1500 random points drawn from a bounding box centered on Minnesota (Latitude: min: 43°N, max: 50°N; Longitude: min: −98°W, max: −89°W). We then used the landscape sample localities to calculate the niche boundaries and density of occurrence for each portion of the range within the environmental PCA space.

For all pairwise comparisons of the range core, early expansion, late expansion, and invasion front, we quantified four measures of niche differentiation. First, we used Schoener’s *D* to quantify the similarity in niche by incorporating both niche breadth and density (Warren et al. 2008). Values of *D* can vary from 0 (no overlap) to 1 (complete overlap). For each pair of regions, we then calculated what proportion of the combined niche space represented niche stability, expansion, and unfilling. In this framework, if ‘A’ represents the older potion of the range (range core) and ‘B’ represents the more recent portion of the range (invasion front), niche stability is the proportion of niche B that overlaps A, niche expansion is the proportion of niche B that does not overlap A, and niche unfilling is the proportion niche A that does not overlap B.

We used permutation tests to determine if values of Schoener’s *D*, expansion, stability, and unfilling were equivalent between the range core, early expansion, late expansion, and invasion front. In the niche equivalency tests, the data were pooled and then randomly assigned to one group of the pairwise range comparisons for 999 permutations. For each permutation, we computed all four statistics. We rejected the null hypothesis of niche equivalency if the observed Schoener’s *D* value was less than 95% of permuted *D* values. Similarly, we rejected niche equivalency based on the combined niche space if observed stability was less than 95% of permuted values and observed niche expansion or unfilling was greater than 95% of permuted values.

#### Ecological niche model

We developed an ecological niche model (ENM) using MaxEnt v.3.4.3 (Merow et al. 2013; Phillips et al. 2017) with the ‘dismo’ package v.1.3-14 in R (Hijmans et al. 2022). Our goal was to identify environmental gradients that could potentially drive phenotypic evolution during range expansion (Elith and Leathwick 2009; Araújo et al. 2019; Morente-López et al. 2022). We built ENMs with the same bioclimatic variables and in the same bounding box as the analyses of niche differentiation. We found that temperature of the warmest quarter (Bio10) had the strongest influence on the probability of occurrence. To examine whether this result was robust, we built three other ENMs using alternative sets of environmental variables (Table S3). In every case, temperature of the warmest quarter was by far the strongest predictor (Table S3).

We built all ENMs using occurrence records from our tissue collection sites and 10,000 background points. We excluded threshold and hinge features during the model building process as the preliminary models that included these features tended to be overspecified. We used five-fold cross-validation to assess model performance; data were randomly partitioned into five equal groupings and 80% of data were used for training and 20% were used for evaluation. Model predictions are a mean of the five cross-validated models.

We evaluated models using AUC and sensitivity, which were calculated using the withheld dataset. AUC characterizes model discrimination ability and ranges between 0 and 1, with higher values indicating greater model performance and a value of 0.5 indicating that model discrimination is no better than random (Phillips and Dudík 2008). Sensitivity quantifies the proportion of correctly identified positives. We calculated sensitivity where the sum of the true positive rate and true negative rate was maximized (threshold = 0.53). We then used the variable permutation importance and percent contribution to identify which environmental variable had the greatest contribution to habitat suitability. We also visualized response curves of each environmental variable to ensure models were not overspecified. Hereafter, “habitat suitability” refers to the probability of occurrence from our ENMs based upon climatic factors.

To test for divergence in germination behavior during range expansion, we focused on mean temperature of the warmest quarter since it most strongly affected predicted habitat suitability in all ENMs. Populations from early in the invasion had high predicted habitat suitability and higher warm season temperatures (all above 20°C) compared to those from later in the invasion, which had lower predicted habitat suitability and lower warm season temperatures (all below 20°C; Fig. 6A,B). Guided by these results, we divided the 14 populations for which we had seed collections into two groups: early (n=8) versus late (n=6) in invasion and exposed them to five temperature regimes (see below).

### Tests for differentiation in germination behavior during range expansion

We collected seeds in 2019 from 14 populations (8 – 24 maternal families/pop; Table S4). Seeds were collected from individuals that were at least 5 meters apart. We stored and after-ripened seeds in an indoor environment for two years prior to the germination experiment (Wicks and Derscheid 1958). Seeds were pooled within populations prior to applying treatments. We were unable to collect seeds from every population used in the population genetic survey because some had already dispersed seeds prior to our collection effort.

We examined the effects of temperature regime and source geographic region (early vs. late in invasion) on germination. Five temperature regimes were designed to mimic the full range of variation in daytime and nighttime temperatures during the spring and summer in this region (14 hour day /10 hour night periods: 15/5 °C, 20/10 °C, 25/15 °C, 30/20 °C, and 35/25 °C). The experiment was conducted in five successive rounds in two growth chambers (Conviron Inc.). We conducted each temperature regime twice – i.e., once in each growth chamber – to control for growth chamber effects.

For each round, we placed ten seeds per population in 60 x 15mm polystyrene Petri dishes containing 2 mL sterile distilled water, lined with one Whatman #1 filter paper, and sealed with parafilm. Each population was replicated three times per chamber for a total of 42 dishes per treatment per chamber or 84 dishes per round. Dishes were wrapped in aluminum foil to block light, which can inhibit germination (Selleck et al., 1962). Every 24 hours we recorded the number of germinated seeds (emergence of radicle) per dish. After each treatment period ended, we tested whether ungerminated seeds were viable by soaking bisected seeds in a 1% Tetrazolium solution for 24 hours (Verma and Majee 2013). Red staining of tissues indicates that seeds are viable. For analyses of germination, we included the number of germinated seeds out of the total number of germinated plus viable (but ungerminated) seeds.

We tested for the effects of temperature regime (categorical), source geographic region, and their interaction on germination probability using a mixed-effects model with a binomial family. Experimental round and population were included as random effects. We evaluated significance with Type III tests. All models were run using the ‘mixed’ function in the ‘afex’ package v.1.3-0 (Singmann et al. 2016) in R v.4.0.2 (R Development Core Team, 2015). We used linear contrasts to test for differences in germination between geographic regions for each temperature regime category individually.

## Results

### Population genomic consequences of range expansion

Analysis of molecular variance (AMOVA) revealed significant partitioning of genetic variance among populations (13.7%; *P* < 0.001), among individuals within populations (8.4%; *P* < 0.001), and within individuals (77.8%; *P* < 0.001) (Table S5). No clonal genotypes were detected in the dataset. STRUCTURE indicated that the optimal number of clusters was three (K = 3) (Table S6). All individuals were assigned primarily to one cluster regardless of where the population was found in the invasive range (i.e., core, early expansion, late expansion, invasion front). There was some evidence of population structure in the other two clusters; however, there was no clear geographic pattern (Figure 2). We also examined K=4, which similarly revealed one major cluster for each individual and a minor one where there was some evidence of population structure. For example, four northern populations near the invasion front share the same minority cluster (Koochiching, St. Louis, Duluth, Aitkin); this cluster is also shared with two populations in the range core (Figure 2).

**Figure 2.**
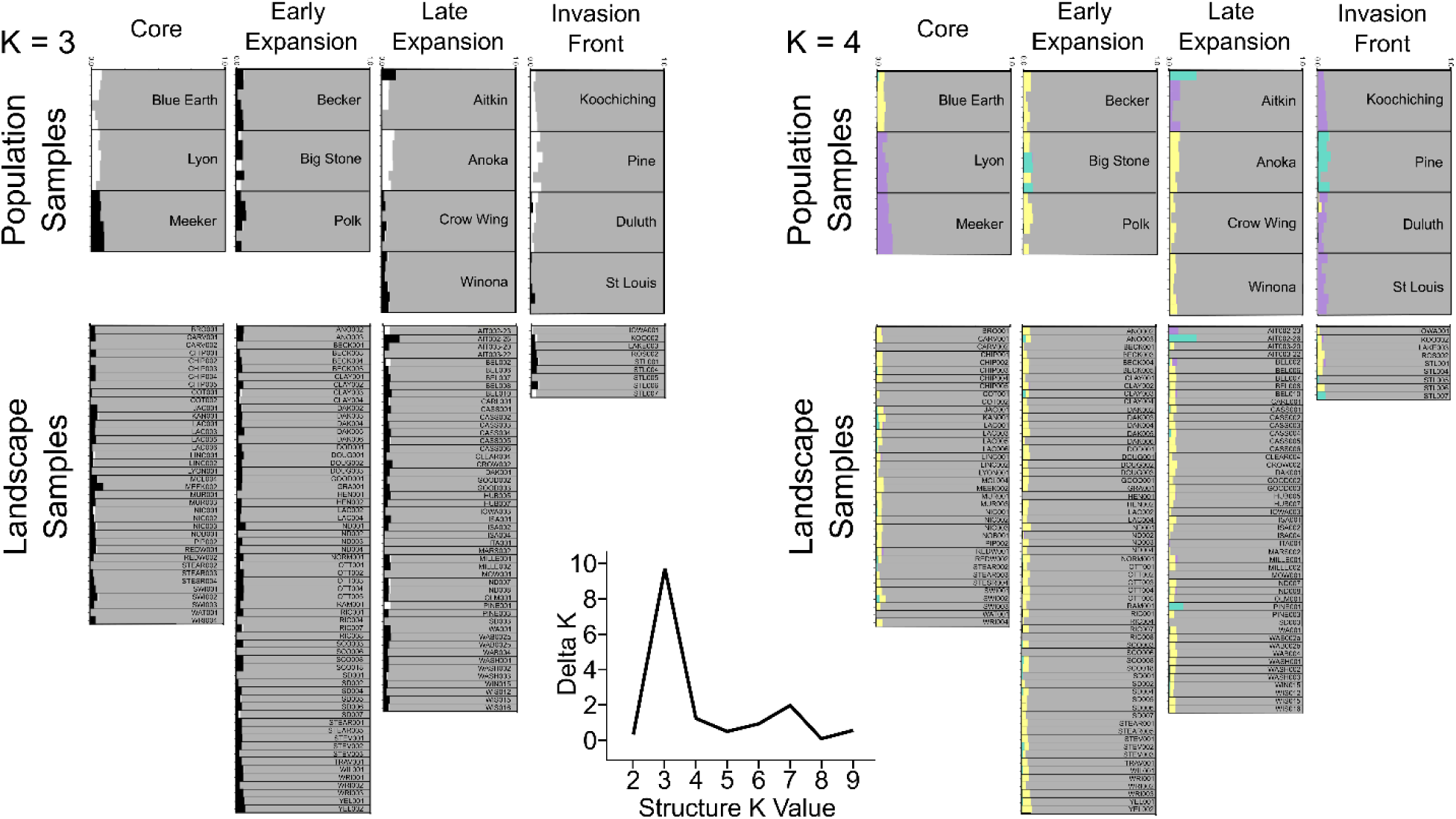
Cluster assignment probability from STRUCTURE analyses (K = 3 and K = 4) for population and landscape samples. Each bar represents one individual, and populations are separated by black lines. A graph of the delta K values supports the optimal number of clusters at K = 3.

The PCA did not indicate substantial population structure across the invaded range based on either population or landscape samples. For the population samples, there was some evidence of differentiation among three populations in or near the range core (Figure 3; PC1 and PC2 explained: 6.8% and 5.9% of variance, respectively). The PCA of landscape samples did not reveal a relationship between genetic similarity and geography over the time course of range expansion (Figure 3; PC1 and PC2 explained 1.7% and 1.3% of variance, respectively).

**Figure 3.**
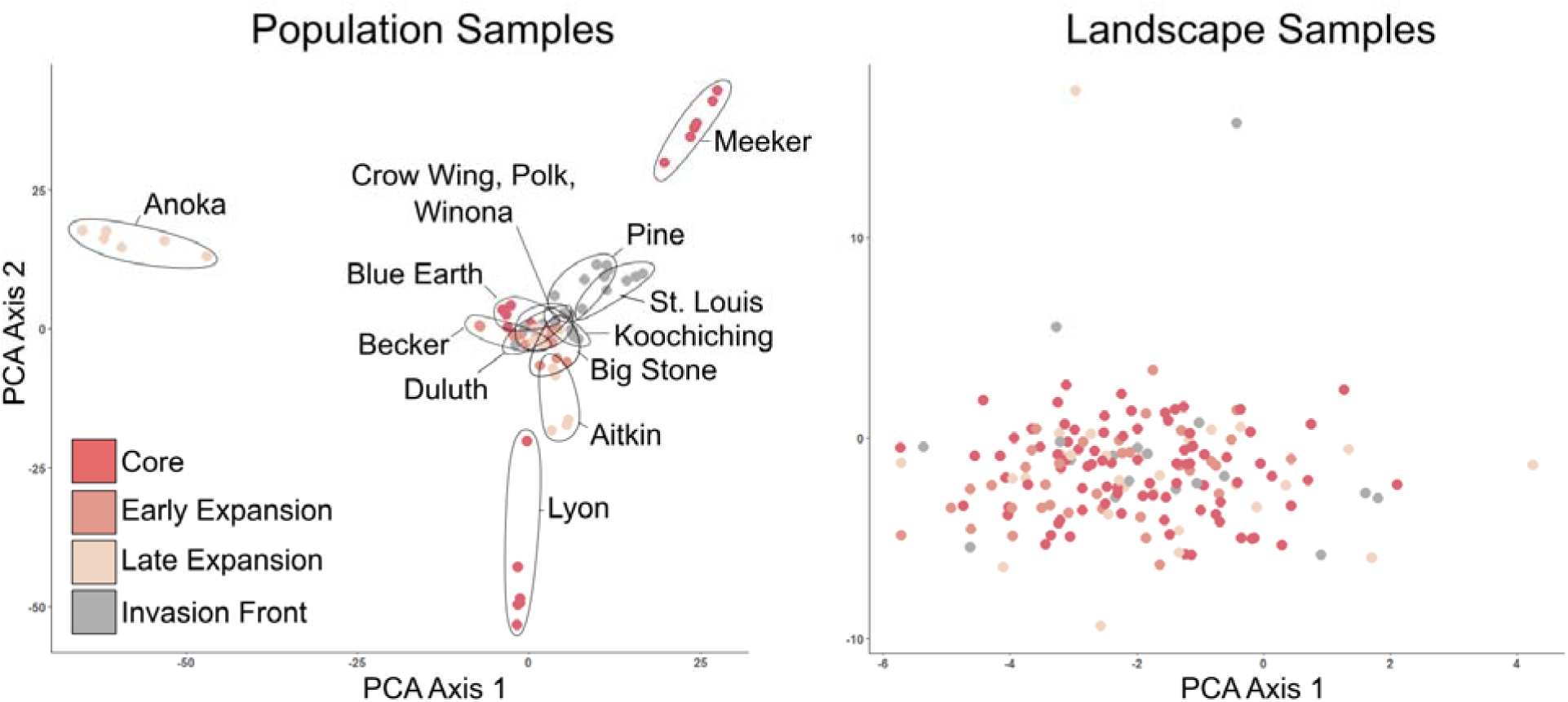
Principal components analysis (PCA) bi-plots for population genomic data from 14 population samples (n = 6 per population) and 157 landscape samples (n = 1 per population).

There was no evidence of isolation by distance (IBD) for either population (*R^2^* = 0.02; *P =* 0.418; Table 2; Table S7) or landscape samples (*R^2^* = 0.05; *P =* 0.676; Table S7). We also did not detect IBD when landscape samples were subset according to the four invasion phases that we defined (Figure 4).

**Figure 4.**
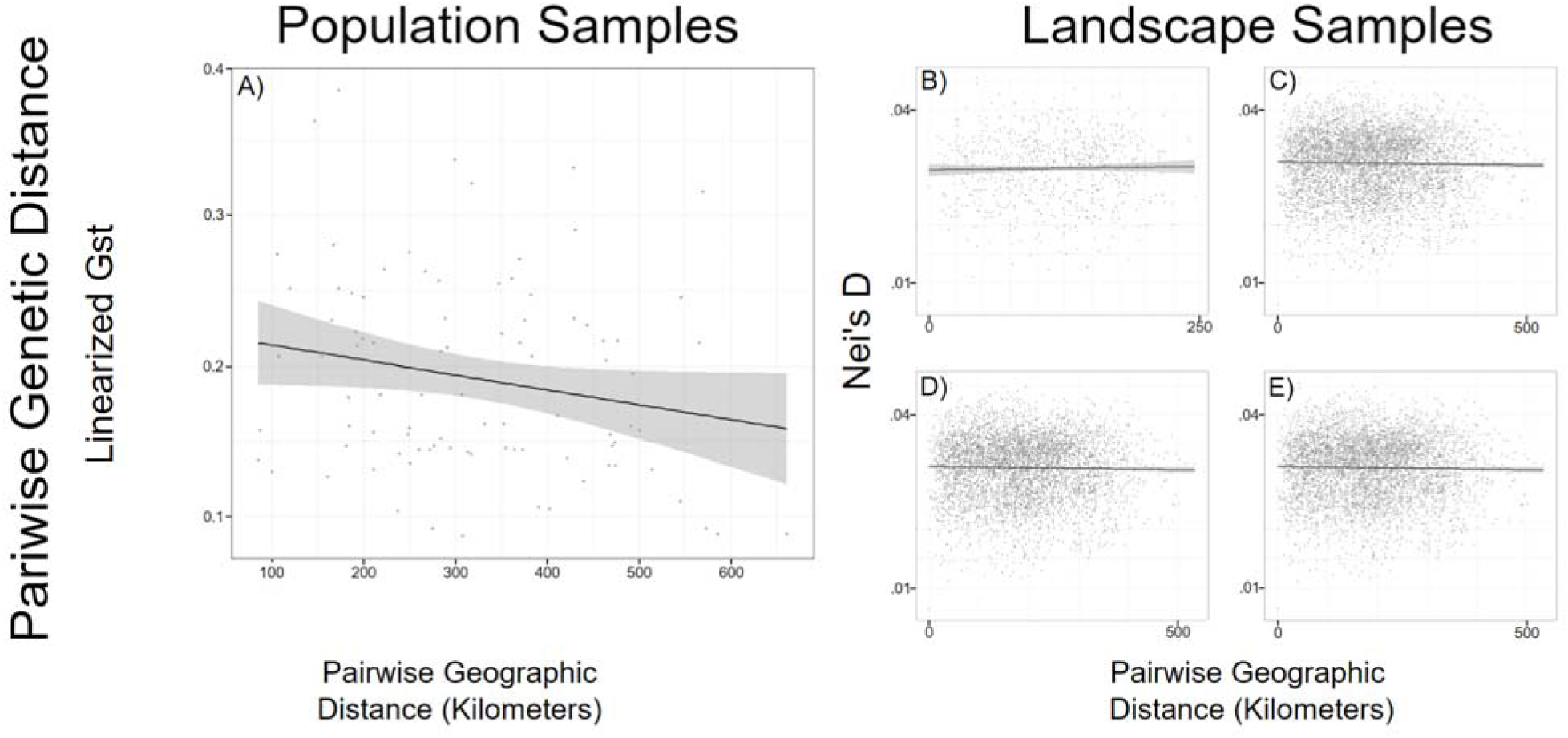
Isolation by distance (IBD) displayed as scatterplots of genetic distance versus geographic distance for A) population samples (A) and B-E) landscape samples (B-E). For landscape samples, we subset individuals for analyses by successive stages of invasion: B) range core (B), C) range core plus early expansion (C), D) range core, early, plus late expansion ranges (D), and E) all samples from across the four phases (E).

**Table 2.**
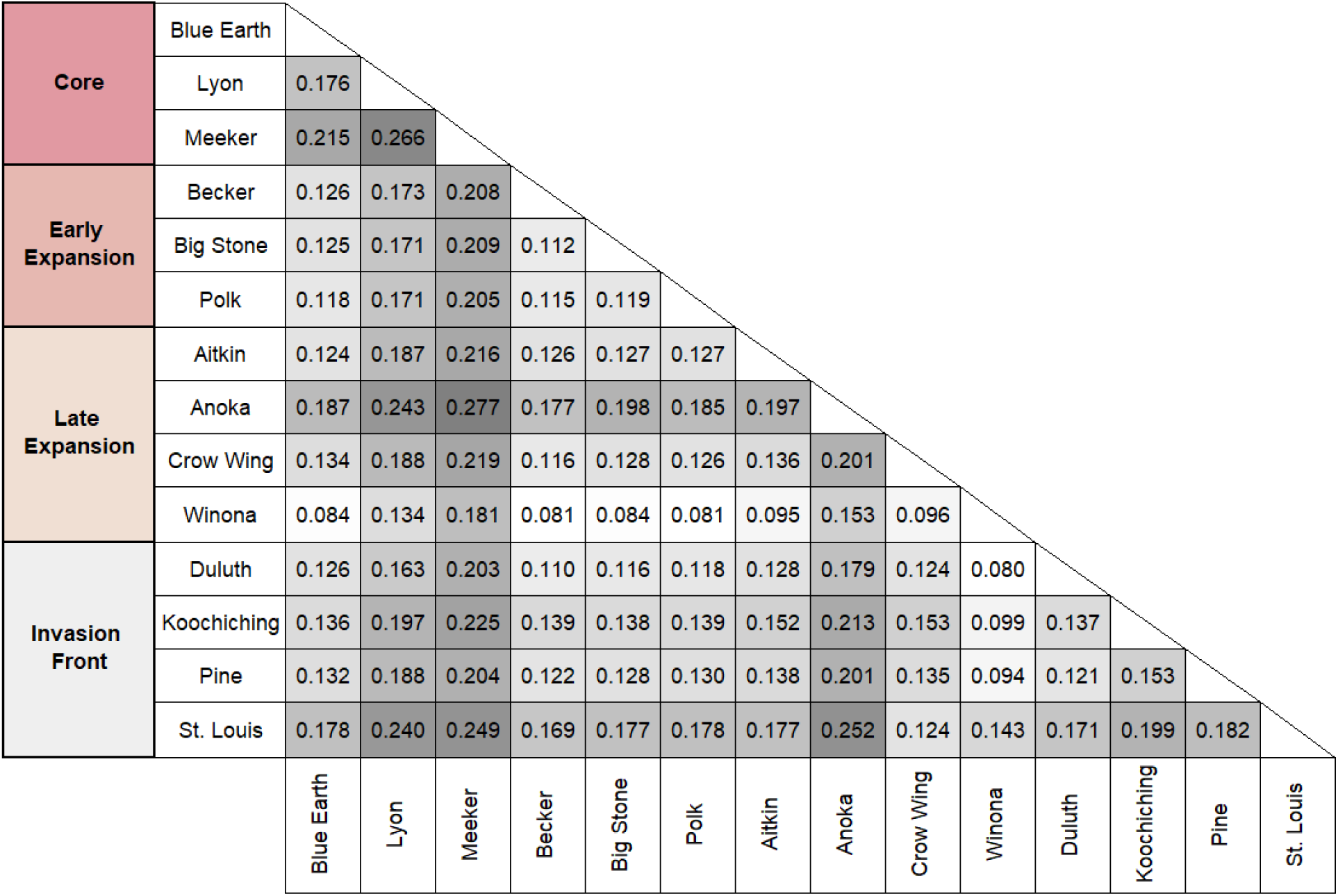
Pairwise estimates of genetic differentiation measured as G_st_ for 14 leafy spurge populations from four phases of range expansion (see Figure 1). Darker versus lighter shading of cells indicates higher versus lower values of pairwise G_st_.

Range expansion from the core area of invasion was accompanied by modest changes in genetic diversity. Heterozygosity declined from the core to the invasion front, as indicated by a significant interaction of latitude and longitude (*P =* 0.013; Table 3). However, the number of private alleles, Tajima’s *D*, and the inbreeding coefficient did not change over the course of range expansion (Table 3). Estimates of N_e_ based on linkage disequilibrium using NeEstimator suggested that N_e_ was large (estimated at infinity) in all but three populations: Blue Earth (N_e_ = 134), Crow Wing (N_e_ = 210), and Koochiching (N_e_ = 17,836). Those three populations occur in the range core, late invasion, and invasion front portions of the range, respectively.

**Table 3.**
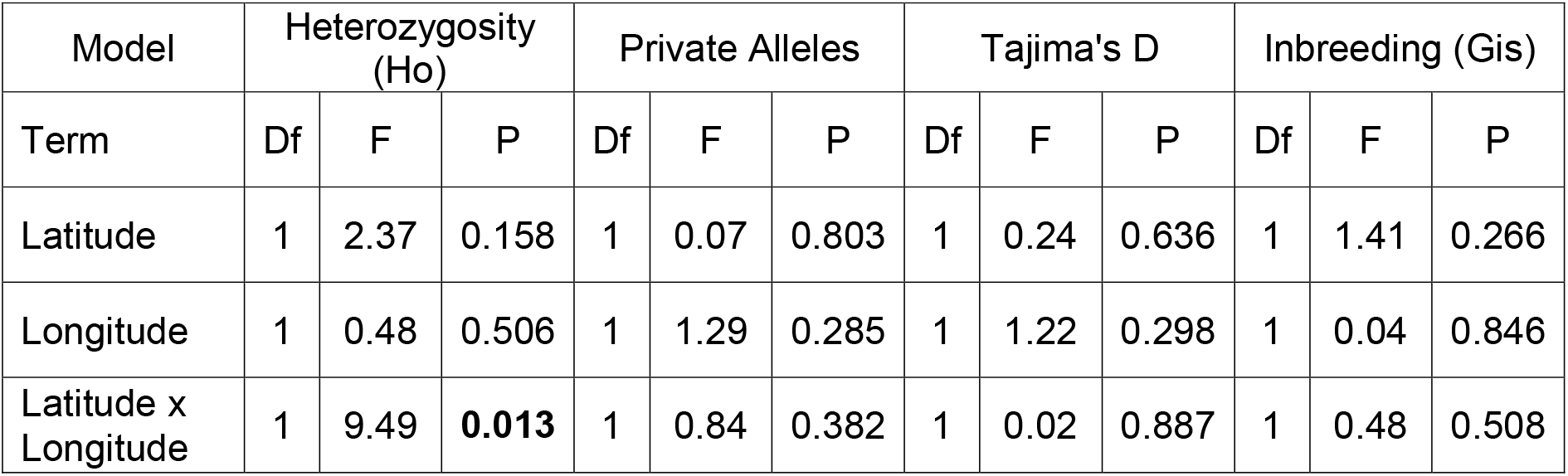
ANOVA testing for a relationship between four metrics of sequence diversity and geography (latitude, longitude, and their interaction) for 14 population samples. Bold indicates p-value less than 0.05.

| Model | Heterozygosity (Ho) |  |  | Private Alleles |  |  | Tajima's D |  |  | Inbreeding (Gis) |  |  |
| --- | --- | --- | --- | --- | --- | --- | --- | --- | --- | --- | --- | --- |
| Term | Df | F | P | Df | F | P | Df | F | P | Df | F | P |
| Latitude | 1 | 2.37 | 0.158 | 1 | 0.07 | 0.803 | 1 | 0.24 | 0.636 | 1 | 1.41 | 0.266 |
| Longitude | 1 | 0.48 | 0.506 | 1 | 1.29 | 0.285 | 1 | 1.22 | 0.298 | 1 | 0.04 | 0.846 |
| Latitude x Longitude | 1 | 9.49 | **0.013** | 1 | 0.84 | 0.382 | 1 | 0.02 | 0.887 | 1 | 0.48 | 0.508 |

### Niche differentiation during range expansion

The climatic niche expanded during invasion. Relative to the range core, the early expansion niche represented a sizable increase in niche breadth (Figure 5A; Table S8; overlap = 0.43; *P* < 0.01; expansion = 0.37; *P* < 0.01; stability = 0.63; *P* < 0.01). When comparing the early expansion and late expansion niches, there was similar evidence for a niche shift (Figure 5B; Table S8; overlap = 0.36, *P* < 0.01; expansion = 0.39, *P* < 0.01; stability = 0.61, *P* < 0.01). Between the late expansion niche and the invasion front, the null hypothesis of niche equivalency was not rejected (Table S8); the invasion front niche was contained within the late expansion niche (Figure 5C). When comparing range core to invasion front, there was near-zero niche overlap (Figure 5D; Table S8; overlap = 0.03, *P* < 0.01; stability = 0.06, *P* < 0.01) and high expansion (Table S8; expansion = 0.94, *P* < 0.01). Niche differences between the core and invasion front were most apparent along environmental axes related to temperature of the warmest quarter and minimum temperature of the coldest month, rather than precipitation (Figure S6).

**Figure 5.**
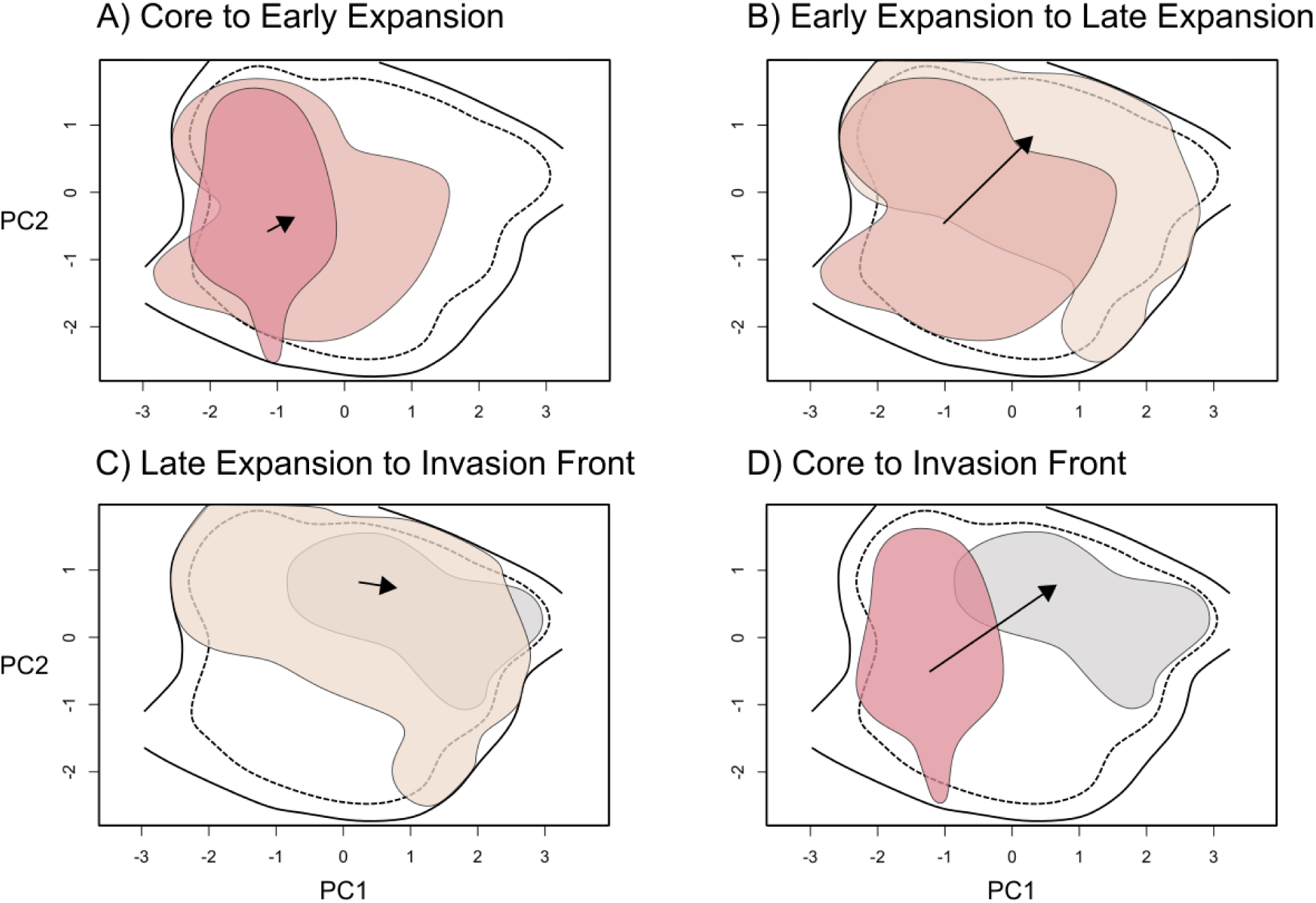
Niche overlap during range expansion in climate niche space. The extent of the background environment is outlined in black (solid = total niche space; dashed = 90% of extent). A) Core (dark pink) versus early expansion (light pink), B) Early expansion (light pink) versus late expansion (beige), C) Late expansion (beige) versus invasion front (grey), D) Core (dark pink) versus invasion front (grey). In all panels, the arrow represents the direction of shift in the centroid of niche space.

#### Ecological niche model

The model AUC (0.79) and sensitivity (0.75) metrics indicated moderately high discrimination and accuracy (Figure 6A). The variable response curves indicated the ENM was not overspecified (Figure S7). The mean temperature of the warmest quarter constituted the most important variable for the predicted probability of occurrence, hereafter “habitat suitability” (percent contribution = 81.5%; permutation importance = 67.8%). Warmer temperatures were associated with an increase in habitat suitability (*R* = 0.67; Figure S8). The minimum temperature of the coldest month had the second highest importance (percent contribution = 11.2%; permutation importance = 17.9%) and was modestly associated with increased habitat suitability (*R* = 0.31; Figure S8). Precipitation of the warmest quarter had the lowest importance (percent contribution = 7.3%; permutation importance = 14.3%) and was weakly correlated with habitat suitability (*R* = 0.02; Figure S8). Populations from early versus late in invasion were distinct along the mean temperature of the warmest quarter axis with a disjunction at 20°C. This temperature also distinguished highly suitable from less suitable habitat in the ENM (Figure 6A & 6B).

**Figure 6.**
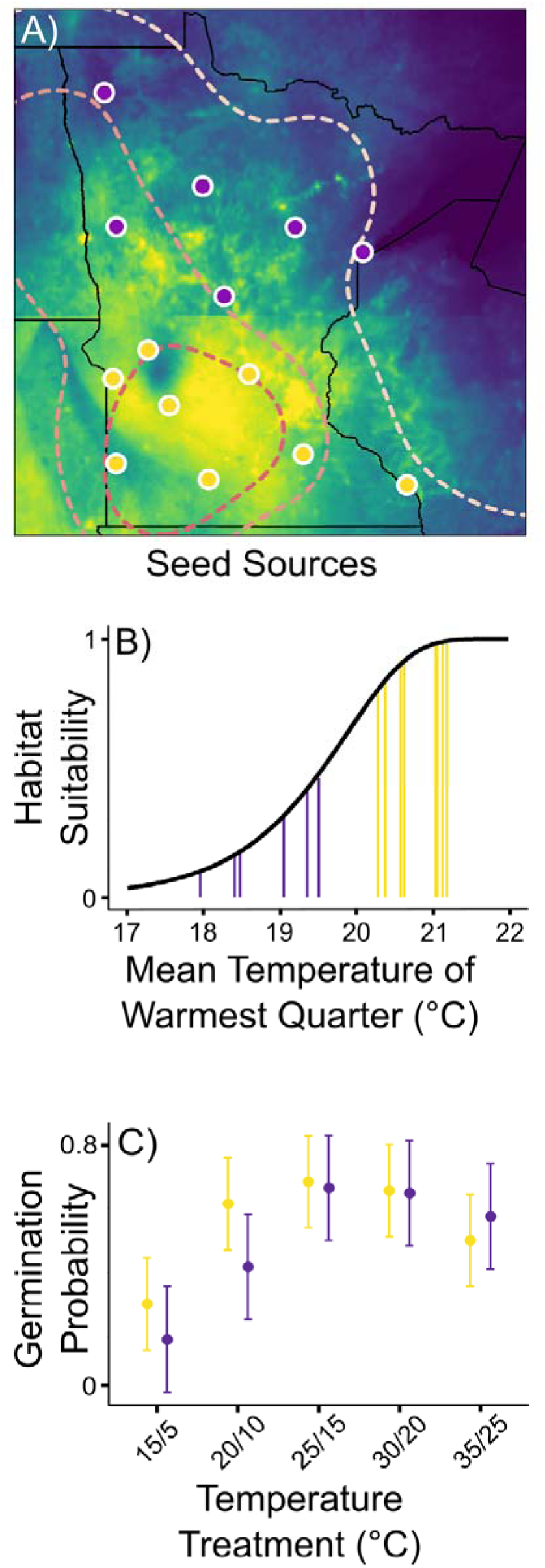
Ecological Niche Model (ENM) habitat suitability projections for seed source populations and germination probability for source regions. A) Habitat suitability projection from leafy spurge ENM. Predicted habitat suitability ranges from 0 (purple) to 1 (yellow). Dashed lines demarcate the chronosequence of range expansion. Seed source populations for the germination experiment are marked and colored yellow (>20°C) or purple (<20°C) depending on their mean temperature of the warmest quarter. B) Variable response curve for the mean temperature of the warmest quarter. Vertical yellow bars correspond to seed source populations from the warmer south and vertical purple bars correspond to source populations from the cooler north. C) Mean germination probability (± SE) by temperature treatment (X axis) for seed source regions. The warmer southern region is shown in yellow and the cooler northern region is shown in purple.

### Evolution of germination behavior during range expansion

Populations from early versus late in invasion responded differently to temperature regimes (source geographic region x temperature regime interaction: *P* = 0.003; Table S9; Figure 6C). The linear contrast between geographic regions for the 20/10 °C treatment, the second coldest treatment, was marginally significant (p = 0.09); contrasts between regions for all other temperature treatments were not significant (p > 0.77; Table S10). Germination probability increased with temperature for both geographic regions and plateaued (*P* < 0.001; Table S9; Figure 6C).

## Discussion

Rapid evolution is increasingly recognized as an important process contributing to range expansion and influencing forecasts of future invasion (Prentis et al. 2008; van Boheemen et al. 2019; Clements and Jones 2021). However, our understanding of the temporal and spatial scale over which niche and trait divergence contribute to invasion at leading range edges remains unresolved. Our study took advantage of a well-documented invasion history to synthesize the consequences of recent range expansion for population genomic diversity, niche breadth, and germination behavior, a trait important in the colonization of new habitats. We found that leafy spurge populations experienced only modest losses in sequence diversity over the chronosequence of invasion and that recruitment within invaded populations occurred via seeds rather than strictly from clonal expansion. Range expansion involved climatic niche expansion and ecological niche models suggested that mean temperature of the warmest quarter had the strongest influence on habitat suitability. Populations differentiated in germination behavior in response to temperature, where leading edge populations exhibited a trend of increased dormancy at low temperatures. Our results suggest that evolution during range expansion may be important to consider in the development of models forecasting range shifts under current and future climates.

Loss of genetic diversity during range expansion may have fitness consequences and limit adaptive capacity (Lee 2002; Dlugosch and Parker 2008). The magnitude of within-population diversity is also potentially important for the success of eradication measures, which may not be effective against all genotypes. We found that range expansion was only accompanied by small losses in heterozygosity but no changes in inbreeding coefficient or the number of private alleles. We also did not observe evidence of genetic bottlenecks based on linkage disequilibrium. We identified some divergence among populations in or near the range core, while others occupied intermediate positions in PCA space. Prevalent long-distance dispersal from the expanding core to the invasion front could reintroduce allelic variation lost due to bottlenecks in the colonization process. Additionally, admixture among independently introduced populations (i.e., invasion fronts not included in our analyses) could contribute to within-population genetic diversity, especially when there is genetic divergence between sources of introductions (Gaskin and Schaal 2002; Keller and Taylor 2010; Gibson et al. 2021). Although we do not observe novel genotypes or elevated numbers of private alleles at the periphery of our focal region, which would be indicative of admixture between invasion fronts, more work is needed to evaluate this potential influence.

Regardless of the mechanism, the extent of within-population variation may have important implications for eradication and management. Although leafy spurge can spread clonally, it does not appear that populations constitute one or few clones that spread widely upon colonization. Instead, recruitment from seeds contributes substantially to population growth. This finding has implications for integrated pest management (IPM) strategies. Management techniques that reduce above-ground productivity and seed production may be important in addition to those that reduce clonal spread by disturbing root systems (e.g. biocontrol agents that feed primarily on roots; Hanson et al. 1997). In addition to natural disturbance, improperly timed grazing or herbicide applications could create open spaces for recruitment from seeds and inadvertently increase density and abundance of the invader (Gaskin et al. 2022). As seeds may remain dormant for as long as five years (Selleck 1962), seed banks may further complicate control efforts. The combination of recruitment from seeds and clonal growth contributes to the population dynamics of leafy spurge, and therefore, multiple eradication approaches may be necessary to cause local population extinction.

In addition to gene flow during invasion, polyploidy may be an important factor influencing losses of genetic variation during colonization bottlenecks and the capacity for range expansion. Polyploids often maintain higher levels of genetic variation (Otto and Whitton 2000) and there is some evidence that phenotypic plasticity is greater in synthetically produced autopolyploids (Mattingly and Hovick 2023). These factors have been used to explain why polyploids may be better invaders than diploids (Pandit et al. 2011). Leafy spurge is an auto-allohexaploid, suggesting that its higher N_e_ should facilitate the maintenance of genetic diversity within populations and minimize divergence among them. Polyploidy has also been suggested to increase the capacity for adaptive evolution (Otto and Whitton 2000), which could also contribute to success as an invader. It is important to recognize that losses of diversity in DNA sequences do not necessarily translate to losses of variation in quantitative traits (Reed and Frankham 2001). Nevertheless, it is possible that polyploidy contributed to rapid invasion in leafy spurge, but more work is needed to distinguish the contribution of polyploidy from other factors.

Accumulating evidence suggests that invasive plant species frequently undergo climatic niche shifts during range expansion (Medley 2010; Atwater et al. 2018; van Boheemen et al. 2019; Bates and Bertelsmeier 2021; but see Petitpierre et al. 2012; Liu et al. 2020). Such niche shifts can complicate the reliability of models that seek to predict future invasions, as these models generally assume niche conservation. Consistent with past findings, we observed climatic niche expansion throughout most of range expansion, except from the late expansion region to the invasion front. From an ENM, we found that warm season temperature had the strongest influence on habitat suitability and therefore may be one source of divergent selection from range core to invasion front. Of course, climate change has already caused poleward shifts in plant species distributions through climatic niche matching (Parmesan 2006; Clements and Ditommaso 2011; Parmesan and Hanley 2015). Therefore, some range expansion may simply involve dispersal to already climatically suitable habitats. However, adaptation may be necessary for continued range expansion, especially where the species is already at its climatic niche limit (Clements and Ditommaso 2011). In leafy spurge, minimal range expansion northward has been observed in the last 30+ years and populations remain small/low density at the invasion front. In addition, ENMs suggest that invasion front populations have very low habitat suitability in terms of climate. Overall, our results are inconsistent with the hypothesis that the leading edge is suitable but expansion is limited by dispersal. Instead, responses to divergent selection may be important for persistence at the leading range edge and for further range expansion. These findings also hold implications for management approaches: recognizing and accounting for niche shifts could improve the accuracy of distribution models for forecasting future invasions, and by extension, help optimize resource allocation for eradication efforts. Integrating knowledge of which environmental factors underly niche expansion will be key to designing more effective management strategies that not only suppress existing populations but also anticipate future invasions.

Divergence in trait expression at a leading range edge can be driven by local adaptation, phenotypic plasticity, and/or maternal environmental effects (Des Roches et al. 2017; Westerband et al. 2021). Phenotypic plasticity is considered important during early stages of invasion because it allows introduced populations to establish in a broader range of environmental conditions (Sexton et al. 2002; Richards et al. 2006; Funk 2008; Lande 2015). In germination traits, plasticity could represent a means of habitat selection and niche construction (Donohue 2003, 2005) by which leafy spurge in the leading edge germinates optimally at the onset of spring conditions (warmer temperatures in northern latitudes). Likewise, rapid evolution during range expansion can result from selection on loci that influence dormancy and/or germination timing (Clements and Ditommaso 2011; Hodgins et al. 2018; Clements and Jones 2021). We found a trend of increased dormancy at lower temperatures in leading edge populations (p = 0.09 for 20/10 °C treatment). One possibility is that germination at colder temperatures exposes seedlings to more unpredictable environments (e.g., late season frost) and thus that selection favored germination later in the season for leafy spurge at its northern range limit. Interestingly, other work has suggested that reduced dormancy evolves at leading range edges (Tabassum and Leishman 2018), contrary to our findings. We cannot exclude the possibility that maternal environmental effects influenced our estimates of germination probability because we used field-collected seeds. However, our findings are contrary to what we would have expected if maternal environmental effects had been important: we would have expected higher germination for leading edge populations in cold treatments because those populations come from colder environments. Although warm season temperature is most strongly associated with habitat suitability and relevant to germination in leafy spurge, it is also possible that other (correlated) variables influenced the evolution of germination behavior. Overall, our results suggest that range expansion involved niche expansion and potentially the evolution of germination timing, which may have been important in establishment at the leading range edge.

Early life history transitions are thought to be under strong selection because of their cascading effects on later life stages (Baskin and Baskin 1971; Marks and Prince 1981; Donohue 2002, 2005). In plant populations, environmental conditions at the time of germination can alter the strength and direction of natural selection on postgermination traits (Donohue et al. 2010; D’Aguillo and Donohue 2023). In turn, this can affect the competitive environment, resource availability, and density-dependent selective agents experienced by populations (Donohue et al. 2010). While we did not investigate postgermination traits, it would be valuable to determine whether germination timing influences performance at later life history stages (e.g., fecundity), especially in leading edge populations. Knowledge of germination timing could help identify optimal windows for applying herbicides or biocontrol agents. If germination timing or success is found to affect performance at later life stages, management strategies could aim to destabilize these windows of opportunity, potentially preventing the establishment of new invasive populations and limiting the growth and persistence of existing ones.

Our results suggest that even over the course of a fairly rapid invasion losses of genomic variation may be minimal. In leafy spurge, this may have occurred because of substantial gene flow during invasion and/or polyploidy. Higher levels of genetic variation can challenge management when genotypes vary in their responses to eradication measures (Gaskin et al. 2020). We also found that trait divergence may have contributed to climatic niche expansion and thus to the spatial extent of invasion. Forecasts of continued invasion typically rely on species distribution models (SDMs), which rarely take into account evolution. As such, models may fail to predict the complete extent of range expansion, or the severity of range infilling. Evolution-free SDMs are likely still valuable for management planning over meaningful spatial and temporal scales in many systems. However, in systems where local adaptation is extensive, forecasts of range shifts with climate change may require the construction of regional SDMs that account for evolution.

## Statements and Declarations

### Author contributions

All authors conceived of the study design, contributed to decisions throughout the analysis process, wrote, and edited the manuscript. T.A.L and R.B.R collected and analyzed data.

### Data Accessibility and Benefit-Sharing

All data will be made publicly available upon publication via the Dryad Digital Repository [DOI].

## Acknowledgments

Funding was provided by the Minnesota Invasive Terrestrial Plants and Pests Center through the Environment and Natural Resources Trust Fund as recommended by the Legislative-Citizen Commission on Minnesota Resources. T.A.L. was supported by the University of Minnesota Doctoral Dissertation Fellowship.The Minnesota Supercomputing Institute provided computing and data storage resources. We thank Yaniv Brandvain and Ken Kozak for discussions and advice on analyses. We also thank Erik Runquist, Brooke Kern, Danielle Schoenecker, Ben Greene, and Isaac Olson for their assistance with data collection.

