## Supplementary Information for "Chronosequence of invasion reveals minimal losses of population genomic diversity, niche expansion, and trait divergence in the polyploid, leafy spurge"

**Supplementary Material**

This document contains supplementary methods, tables, and figures for the manuscript titled: *“Chronosequence of invasion reveals minimal losses of population genomic diversity, niche expansion, and trait divergence in the polyploid, leafy spurge”.*

Table of Contents

Section 1: Supplementary methods

Table S1. Sampling information and localities for leafy spurge population and individual samples.

Table S2. Tests for genetic bottlenecks based on linkage disequilibrium with NeEstimator.

Table S3. Alternative sets of bioclimatic variables tested in ENMs.

Table S4. Seed population localities and environmental details for germination experiment.

Table S5. Analyses of Molecular Variance (AMOVA) results.

Table S6. Analyses of population structure with STRUCTURE with the Evanno Delta K method.

Table S7. Analyses of isolation by distance and Mantel tests.

Table S8. Niche differentiation permutation tests.

Table S9. Analysis of variance table for mixed effect models on germination rate.

Table S10. Tests for general linear contrasts.

Figure S1. Point map of leafy spurge ca. 1933 reproduced from Hanson and Rudd (1933).

Figure S2. Filtering by minor allele frequency to remove homoeologous alleles.

Figure S3. Residuals versus fitted plot for private alleles among populations.

Figure S4. PCA of 19 BIOCLIM variables.

Figure S5. Eigenvectors of environmental variables used in niche analyses and ENM.

Figure S6. Axes of environmental variables used in niche analyses between core and invasion front.

Figure S7. Variable response curves for the ecological niche model with locations of seed sources.

Figure S8. Correlation between habitat suitability and environmental variables.

**Section 1: Supplementary methods**

Genotyping by Sequencing (GBS) data for all samples were initially delivered by the University of Minnesota Genomics Center to the Minnesota Supercomputing Institute in the .fastq file format. We first removed GBS adapter sequences using custom Perl scripts provided by the UMGC (<https://bitbucket.org/jgarbe/gbstrim/src/master/>).

We next used the Trimmomatic software (Bolger, Lohse, & Usadel, 2014) to remove reads with an average Phred quality score < 30 in a 4 bp sliding window, discard leading and trailing bases with a quality score < 30, and crop all reads to 80 bp in length. We next used Stacks v.2.5.9 (Rochette et al. 2019) to build loci *de novo* (i.e., without aligning reads to a reference genome). Stacks consists of several scripts that are executed sequentially (i.e. *ustacks, cstacks, sstacks, gstacks)* with the *denovo_map.pl* Perl script. We examined several key parameters in the Stacks pipeline: -m (the minimum number of identical raw reads required to form a putative allele), -M (the number of mismatches allowed between alleles to form a locus), and -n (the number of mismatches allowed between loci during construction of the catalog) to optimize the number of SNPs, assembled loci, and polymorphic loci (Paris et al. 2017). We found the parameter combination of (m = 3, M = 4, n = 3) was an optimum to maximize these criteria. Because leafy spurge is an auto-allohexaploid species (Horvath et al. 2018; Schulz-Schaeffer and Gerhardt), we specified “*-X ustacks –max_locus_stacks 7”* while running the *denovo_map.pl* pipeline to allow the possible number of alleles per loci to reach seven instead of the default three (Dufresne et al. 2014). However, as the Stacks *populations* script calls genotypes assuming a diploid model, we instead used the Bayesian approach polyRAD designed for variant calling in polyploid organisms.

Briefly, we applied the polyRAD function *read_stacks* to import read depth information from the catalog and matches files output by Stacks *cstacks* and *sstacks* functions, respectively. In polyRAD, we specified *ploidy = 6* and filtered loci by setting the options *min.ind.with.minor.alleles = 7* to specify the minimum number of individuals with reads in a minor allele needed to retain a locus and *min.ind.with.reads = 172* to require that a minimum of 70% (172/247 ~ 70%) of individuals with reads needed were needed to retain a locus.

We calculated the H_ind_/H_e_ statistic in polyRAD as a ratio of the individual heterozygosity to expected heterozygosity (Clark et al. 2022). We calculated and plotted the H_ind_/H_e_ statistic by locus to evaluate marker quality and to assess whether a marker represents one Mendelian locus or multiple collapsed paralogous loci, based upon read depth distribution in a population (Clark et al. 2022). We identified a peak in the H_ind_/H_e_ plot at 0.28, indicating well-behaved markers (i.e., markers likely following a pattern of Mendelian inheritance). We next applied the *InbreedingFromHindHe* function to estimate inbreeding and to simulate the expected distribution of H_ind_/H_e_ under the assumption that all markers were behaving in a Mendelian fashion. Based on this simulated distribution, we restricted the minimum and maximum H_ind_/H_e_ thresholds to capture 95% of the samples [0.050 - 0.353], then filtered markers where the H_ind_/H_e_ statistic were outside of this range. Using these filtered markers, we next performed permutation tests for overdispersion (i.e., the observation that variation is higher than would be expected) to examine how the read depth distribution deviates from expectation under a binomial (i.e. diploid) distribution. We set the parameter to 8 (with tests ranging from 2 to 50) to minimize the amount of overdispersion present in the markers through visual inspection. Using these parameters, we used the polyRAD function *IterateHWE* to estimate the genotype posterior probabilities without a prior assumption of population structure. For each individual at each locus, we exported the most probable genotype for subsequent analyses.

Table S1. Population name, geographic location, and number of individuals sampled per population, totaling 14 population samples and 157 landscape samples of leafy spurge. Population samples are indicated in bold.

| Population | Latitude | Longitude | Number of Individuals Sampled |
| --- | --- | --- | --- |
| AIT002-23 | 46.85855 | -93.61362 | 1 |
| AIT002-26 | 46.97621 | -93.72486 | 1 |
| **AIT002** | **46.98386** | **-93.71759** | **6** |
| AIT003-20 | 46.99886 | -93.32183 | 1 |
| AIT003-22 | 46.73479 | -93.27241 | 1 |
| **ANO001** | **45.28849** | **-93.12503** | **6** |
| ANO002 | 45.20712 | -93.29607 | 1 |
| ANO003 | 45.11867 | -93.23818 | 1 |
| BECK001 | 46.85924 | -95.87857 | 1 |
| **BECK002** | **46.87731** | **-96.05467** | **6** |
| BECK003 | 46.80351 | -96.14682 | 1 |
| BECK004 | 46.67106 | -95.69111 | 1 |
| BECK005 | 46.68252 | -95.68796 | 1 |
| BEL002 | 47.87728 | -94.28244 | 1 |
| BEL006 | 48.14117 | -95.25442 | 1 |
| BEL007 | 48.23014 | -95.23993 | 1 |
| BEL008 | 48.34406 | -95.12022 | 1 |
| BEL010 | 47.46123 | -94.85712 | 1 |
| **BIG001** | **45.51940** | **-96.55387** | **6** |
| **BLU001** | **44.15582** | **-94.08855** | **6** |
| BRO001 | 44.32644 | -94.52117 | 1 |
| CARL001 | 46.48228 | -92.72164 | 1 |
| CARV001 | 44.87955 | -93.68686 | 1 |
| CARV002 | 44.80925 | -93.55538 | 1 |
| CASS001 | 47.41126 | -94.07930 | 1 |
| CASS002 | 47.37376 | -94.54605 | 1 |
| CASS003 | 46.98354 | -94.52289 | 1 |
| CASS004 | 46.91974 | -94.68527 | 1 |
| CASS005 | 47.05462 | -94.60547 | 1 |
| CASS006 | 47.05519 | -94.48992 | 1 |
| CHIP001 | 45.14534 | -96.02128 | 1 |
| CHIP002 | 45.11188 | -95.86667 | 1 |
| CHIP003 | 44.99849 | -95.75196 | 1 |
| CHIP004 | 45.08466 | -95.56668 | 1 |
| CHIP005 | 44.93980 | -95.41192 | 1 |
| CLAY001 | 46.75890 | -96.36533 | 1 |
| CLAY002 | 46.48890 | -96.72008 | 1 |
| CLAY003 | 47.06523 | -96.43211 | 1 |
| CLAY004 | 47.00628 | -96.28663 | 1 |
| CLEAR004 | 47.14589 | -95.55986 | 1 |
| COT001 | 43.93810 | -94.92046 | 1 |
| COT002 | 43.73148 | -95.06834 | 1 |
| **CROW001** | **46.38368** | **-94.22386** | **6** |
| CROW002 | 46.22316 | -94.35923 | 1 |
| DAK001 | 44.70575 | -92.76531 | 1 |
| DAK002 | 44.73876 | -93.01722 | 1 |
| DAK003 | 44.76540 | -93.19692 | 1 |
| DAK004 | 44.76735 | -92.93372 | 1 |
| DAK005 | 44.67342 | -92.81342 | 1 |
| DAK006 | 44.52157 | -92.93000 | 1 |
| DOD001 | 44.05221 | -92.99615 | 1 |
| DOUG001 | 45.94085 | -95.35931 | 1 |
| DOUG003 | 45.83126 | -95.69664 | 1 |
| GOOD001 | 44.27789 | -92.93795 | 1 |
| GOOD002 | 44.61753 | -92.74396 | 1 |
| GOOD003 | 44.56307 | -92.56240 | 1 |
| GRA001 | 45.90815 | -95.87166 | 1 |
| HEN001 | 45.02023 | -93.32475 | 1 |
| HEN002 | 44.90440 | -93.19830 | 1 |
| HUB005 | 46.85735 | -94.72389 | 1 |
| HUB007 | 46.82788 | -94.89485 | 1 |
| IOWA001 | 42.06415 | -93.56893 | 1 |
| IOWA003 | 43.33455 | -93.34914 | 1 |
| ISA001 | 45.52978 | -93.36012 | 1 |
| ISA002 | 45.54482 | -93.23291 | 1 |
| ISA004 | 45.50387 | -93.02557 | 1 |
| ITA001 | 47.75381 | -94.26960 | 1 |
| JAC001 | 43.80981 | -95.20918 | 1 |
| KAN001 | 45.33318 | -95.05110 | 1 |
| **KOO001** | **48.60145** | **-93.39640** | **6** |
| KOO002 | 48.34507 | -93.69505 | 1 |
| LAC001 | 44.84817 | -96.29217 | 1 |
| LAC002 | 45.07167 | -96.39440 | 1 |
| LAC003 | 45.02712 | -96.19628 | 1 |
| LAC004 | 45.21348 | -96.16614 | 1 |
| LAC005 | 45.11800 | -96.11027 | 1 |
| LAC006 | 44.83915 | -96.09917 | 1 |
| LAKE003 | 47.10945 | -91.52165 | 1 |
| LINC001 | 44.25168 | -96.28914 | 1 |
| LINC002 | 44.21195 | -96.30483 | 1 |
| LYON001 | 44.35820 | -95.92827 | 1 |
| **LYON002** | **44.33427** | **-95.81576** | **6** |
| MARS002 | 48.50045 | -96.48752 | 1 |
| MCL004 | 44.92002 | -94.46786 | 1 |
| **MEEK001** | **44.93534** | **-94.63685** | **6** |
| MEEK002 | 45.31159 | -94.49233 | 1 |
| MILLE001 | 46.12822 | -93.67675 | 1 |
| MILLE002 | 46.33261 | -93.84686 | 1 |
| MOW001 | 43.72490 | -92.74960 | 1 |
| MUR001 | 44.36306 | -95.48021 | 1 |
| MUR003 | 43.99333 | -96.04376 | 1 |
| ND001 | 46.57281 | -96.82062 | 1 |
| ND002 | 46.87634 | -96.84367 | 1 |
| ND003 | 47.20133 | -96.85727 | 1 |
| ND004 | 47.91583 | -97.09736 | 1 |
| ND007 | 46.89406 | -98.90747 | 1 |
| ND008 | 46.12948 | -98.57290 | 1 |
| NIC001 | 44.19654 | -94.05753 | 1 |
| NIC002 | 44.31193 | -94.01828 | 1 |
| NIC003 | 44.29227 | -94.18727 | 1 |
| NOB001 | 43.81974 | -95.60274 | 1 |
| NORM001 | 47.29587 | -96.53400 | 1 |
| OLM001 | 44.10802 | -92.52961 | 1 |
| OTT001 | 46.23161 | -96.03605 | 1 |
| OTT002 | 46.31913 | -96.13459 | 1 |
| OTT003 | 46.31803 | -95.67730 | 1 |
| OTT004 | 46.53053 | -95.97296 | 1 |
| OTT005 | 46.67634 | -95.96223 | 1 |
| PINE001 | 45.98769 | -92.53716 | 1 |
| **PINE002** | **46.04031** | **-92.35830** | **6** |
| PINE003 | 46.32158 | -92.79186 | 1 |
| PIP002 | 43.89599 | -96.36810 | 1 |
| **POLK002** | **47.75272** | **-96.25486** | **6** |
| RAM001 | 45.04734 | -93.22468 | 1 |
| REDW001 | 44.23877 | -95.21953 | 1 |
| REDW002 | 44.22671 | -95.37033 | 1 |
| RIC001 | 44.30155 | -93.18172 | 1 |
| RIC004 | 44.32706 | -93.16534 | 1 |
| RIC007 | 44.36140 | -93.19106 | 1 |
| RIC008 | 44.44749 | -93.16923 | 1 |
| ROS002 | 48.84372 | -95.01659 | 1 |
| SCO003 | 44.57367 | -93.31000 | 1 |
| SCO006 | 44.64500 | -93.30224 | 1 |
| SCO008 | 44.68170 | -93.34550 | 1 |
| SCO018 | 44.77142 | -93.39660 | 1 |
| SD001 | 43.50576 | -96.74930 | 1 |
| SD002 | 43.50773 | -96.75977 | 1 |
| SD003 | 43.62370 | -97.08950 | 1 |
| SD004 | 43.55347 | -96.72285 | 1 |
| SD005 | 44.89197 | -97.06898 | 1 |
| SD006 | 45.10679 | -96.62681 | 1 |
| SD007 | 45.25535 | -96.34096 | 1 |
| STEAR001a | 45.48404 | -94.13496 | 1 |
| STEAR001b | 45.52225 | -94.15565 | 1 |
| STEAR002 | 45.47006 | -94.77898 | 1 |
| STEAR003a | 45.57121 | -95.06326 | 1 |
| STEAR003b | 45.57087 | -94.94727 | 1 |
| STEV001 | 45.58825 | -95.75842 | 1 |
| STEV002 | 45.74536 | -96.00158 | 1 |
| STEV003 | 45.47120 | -96.12992 | 1 |
| STL001 | 46.71458 | -92.20389 | 1 |
| **STL003** | **46.76025** | **-92.10583** | **6** |
| STL004 | 46.77430 | -92.14129 | 1 |
| STL005 | 46.81632 | -92.10033 | 1 |
| STL006 | 47.74023 | -91.48627 | 1 |
| STL007 | 46.83622 | -92.01415 | 1 |
| **STL008** | **47.71546** | **-91.97007** | **6** |
| SWI001 | 45.36108 | -95.37393 | 1 |
| SWI002 | 45.35432 | -95.49200 | 1 |
| SWI003 | 45.29152 | -95.63898 | 1 |
| TRAV001 | 45.76255 | -95.61070 | 1 |
| WA001 | 44.27861 | -91.93290 | 1 |
| WAB002a | 44.28808 | -92.46506 | 1 |
| WAB002b | 44.28854 | -92.45205 | 1 |
| WAB003 | 44.28829 | -92.46344 | 1 |
| WASH001 | 44.84504 | -92.79242 | 1 |
| WASH002 | 45.03924 | -92.79358 | 1 |
| WASH003 | 45.07217 | -92.85461 | 1 |
| WAT001 | 44.06202 | -94.75903 | 1 |
| WIL001 | 46.25828 | -96.57448 | 1 |
| **WIN011** | **44.03631** | **-91.61718** | **6** |
| WIN015 | 43.99584 | -91.47315 | 1 |
| WIS012 | 43.95083 | -90.77962 | 1 |
| WIS015 | 43.78708 | -91.07110 | 1 |
| WIS016 | 43.79511 | -91.18027 | 1 |
| WRI001 | 45.29499 | -93.76627 | 1 |
| WRI002 | 45.30602 | -94.09112 | 1 |
| WRI003 | 45.30265 | -93.94783 | 1 |
| WRI004 | 45.08177 | -93.82798 | 1 |
| YEL001 | 44.77654 | -96.44941 | 1 |
| YEL002 | 44.79028 | -96.37692 | 1 |

Table S2. Tests for genetic bottlenecks based on linkage disequilibrium with NeEstimator. Estimated effective population size (Ne^) values shown for each population sample.

| **Range Position** | **Population Genetic Samples** | **Latitude** | **Longitude** | **Ne^** |
| --- | --- | --- | --- | --- |
| **Core** | Blue Earth | 44.16 | -94.09 | 134.40 |
|  | Lyon | 44.33 | -95.82 | Infinite |
|  | Meeker | 44.94 | -94.64 | Infinite |
| **Early Expansion** | Becker | 46.88 | -96.05 | Infinite |
|  | Big Stone | 45.52 | -96.55 | Infinite |
|  | Polk | 47.75 | -96.25 | Infinite |
| **Late Expansion** | Aitkin | 46.98 | -93.72 | Infinite |
|  | Anoka | 45.29 | -93.13 | Infinite |
|  | Crow Wing | 46.38 | -94.22 | 210.30 |
|  | Winona | 44.04 | -91.62 | Infinite |
| **Invasion Front** | Duluth | 46.76 | -92.11 | Infinite |
|  | Koochiching | 48.60 | -93.40 | 17835.60 |
|  | Pine | 46.04 | -92.36 | Infinite |
|  | St. Louis | 47.72 | -91.97 | Infinite |

Table S3. Alternative bioclimatic variables tested in ENMs, with each model’s bioclimatic variable percent contribution and importance.

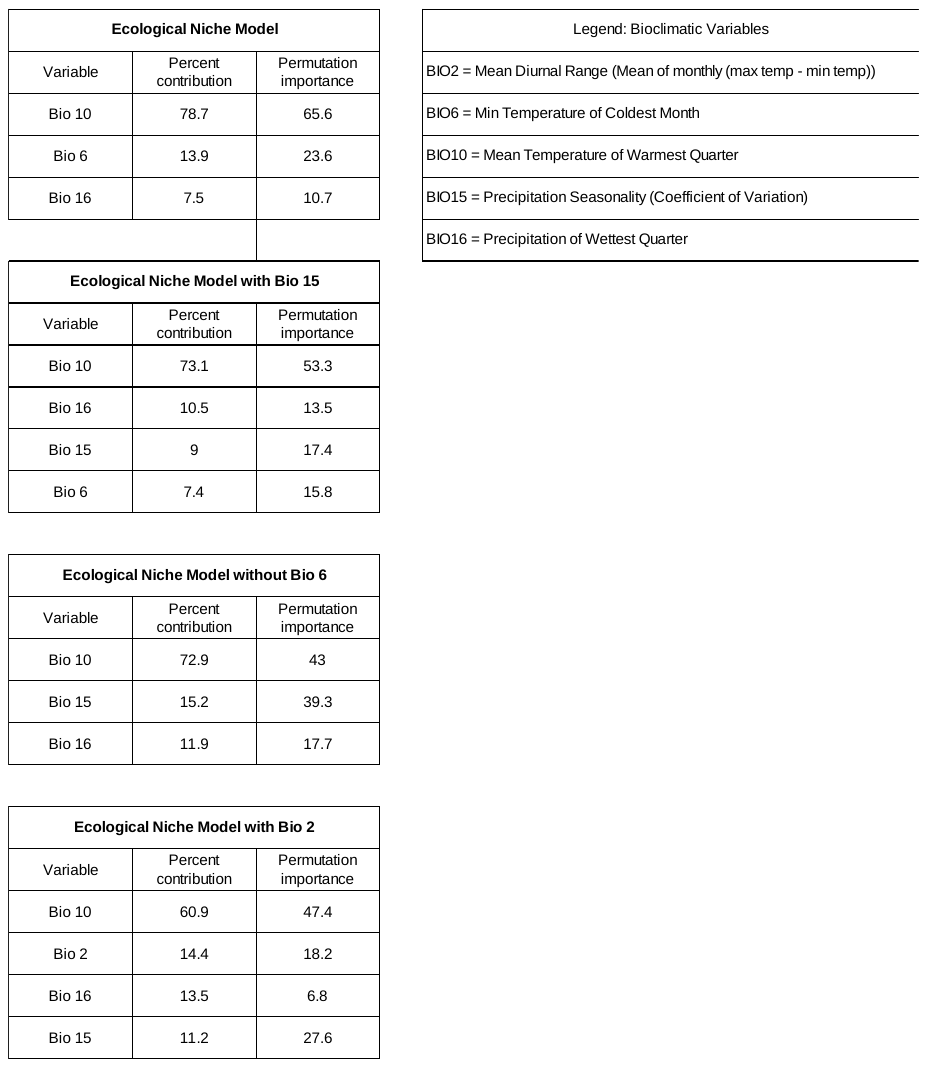

Table S4. Leafy spurge seed source localities as latitude and longitude coordinates. Populations are categorized by range position, and include information about the predicted habitat suitability from the ecological niche model, as well as specific environmental variables at each site.

| Population | Latitude | Longitude | Range Position | Environment | Habitat Suitability | Mean Temp Warm Q | Min Temp Cold Q | Precip Warm Q |
| --- | --- | --- | --- | --- | --- | --- | --- | --- |
| MARS002 | 48.50 | -96.49 | Late Expansion | Marginal | 0.14 | 18.48 | -22.60 | 222.98 |
| BEL010 | 47.46 | -94.86 | Late Expansion | Marginal | 0.52 | 19.05 | -21.80 | 281.75 |
| AIT003 | 47.00 | -93.32 | Late Expansion | Marginal | 0.43 | 17.97 | -20.80 | 323.79 |
| CLAY004 | 47.01 | -96.29 | Early Expansion | Optimal | 0.62 | 19.52 | -20.40 | 257.05 |
| MOR001 | 46.21 | -94.50 | Late Expansion | Optimal | 0.58 | 19.36 | -19.20 | 282.10 |
| STL001 | 46.71 | -92.20 | Invasion Front | Marginal | 0.21 | 18.42 | -18.05 | 305.53 |
| STEV001 | 45.59 | -95.76 | Early Expansion | Optimal | 0.62 | 20.39 | -18.00 | 242.23 |
| WRI002 | 45.31 | -94.09 | Late Expansion | Optimal | 0.87 | 20.63 | -17.50 | 296.87 |
| SD007 | 45.26 | -96.34 | Early Expansion | Optimal | 0.84 | 21.05 | -17.41 | 250.15 |
| LINC001 | 44.25 | -96.29 | Core | Optimal | 0.53 | 20.28 | -17.18 | 255.40 |
| CHIP005 | 44.94 | -95.41 | Core | Optimal | 0.81 | 21.13 | -17.00 | 249.78 |
| ANO002 | 45.21 | -93.30 | Early Expansion | Optimal | 0.93 | 21.20 | -16.73 | 315.31 |
| RIC007 | 44.36 | -93.19 | Early Expansion | Optimal | 0.67 | 20.58 | -16.41 | 308.81 |
| WAT001 | 44.06 | -94.76 | Core | Optimal | 0.75 | 21.20 | -15.90 | 281.07 |
| WIN015 | 44.00 | -91.47 | Late Expansion | Optimal | 0.54 | 21.06 | -15.04 | 308.90 |

Table S5. Analysis of Molecular Variance (AMOVA) tables demonstrating differences in partitioning of genetic variance among populations, among individuals within populations, and within individuals. Significant P-values for AMOVA permutation tests are indicated with bolded terms.

| Results | Df | Sum Sq | Mean Sq |
| --- | --- | --- | --- |
| Between Populations | 13 | 41059.66 | 3158.4356 |
| Between Samples Within Populations | 70 | 49110.2 | 564.485 |
| Within Samples | 420 | 172924 | 342.4238 |
| Total | 503 | 263093.86 | 434.8659 |
| Components of Covariance |  | Sigma | % |
| Variations Between Populations |  | 60.38 | 13.73 |
| Variations Between Samples Within Populations |  | 37.01 | 8.41 |
| Variations Within Samples |  | 342.42 | 77.86 |
| Total Variance |  | 439.81 | 100 |
| Phi Statistic |  |  | Phi |
| Phi Samples Total |  |  | 0.2214 |
| Phi Samples Population |  |  | 0.0975 |
| Phi Samples Population-Total |  |  | 0.1239 |

| Test | Obs | Std.Obs | Alternative | Pvalue |
| --- | --- | --- | --- | --- |
| Variation within samples | 342.423 | -34.549 | less | **0.001** |
| Variation between samples | 37.01 | 10.229 | greater | **0.001** |
| Variation between populations | 60.387 | 53.873 | greater | **0.001** |

Table S6. Results of population structure analyses with STRUCTURE and the Evanno Delta K method. The optimum population assignment of K=3 is demonstrated by the maximum Delta K value as bolded term.

| # This document produced by structureHarvester.py v0.6.94 July 2014 core vA.2 July 2014 | | | | | | |
| --- | --- | --- | --- | --- | --- | --- |
| # http://www.structureharvester.com/ | | | |  |  |  |
| # https://github.com/dentearl/structureHarvester/ | | | | |  |  |
| # http://taylor0.biology.ucla.edu/structureHarvester | | | | | |  |
| # http://users.soe.ucsc.edu/~dearl/software/structureHarvester | | | | | | |
| # Written by Dent Earl, dearl (a) soe ucsc edu. | | | | |  |  |
| # CITATION: | |  |  |  |  |  |
| # Earl, Dent A. and vonHoldt, Bridgett M. (2012) | | | | |  |  |
| # STRUCTURE HARVESTER: a website and program for visualizing | | | | | | |
| # STRUCTURE output and implementing the Evanno method. | | | | | |  |
| # Conservation Genetics Resources 4(2) 359-361. DOI: 10.1007/s12686-011-9548-7 | | | | | | |
| # Stand-alone version: v0.6.94 July 2014 | | | |  |  |  |
| # Core version: vA.2 July 2014 | | |  |  |  |  |
| # File generated at 2022-Jun-25 00:28:54 CDT | | | | |  |  |
| # |  |  |  |  |  |  |
| ########## | |  |  |  |  |  |
| # K | Reps | Mean LnP(K) | Stdev LnP(K) | Ln'(K) | \|Ln''(K)\| | Delta K |
| 1 | 10 | -1006879 | 0.854 | NA | NA | NA |
| 2 | 10 | -1005283 | 12453.87 | 1595.94 | 4121.79 | 0.330965 |
| **3** | **10** | **-999565** | **355.8498** | **5717.73** | **3464.31** | **9.735315** |
| 4 | 10 | -997311 | 1093.529 | 2253.42 | 1364.64 | 1.247924 |
| 5 | 10 | -996423 | 2681.898 | 888.78 | 1321.9 | 0.492897 |
| 6 | 10 | -996856 | 2430.101 | -433.12 | 2290.47 | 0.942541 |
| 7 | 10 | -994998 | 1598.637 | 1857.35 | 3110.44 | 1.945683 |
| 8 | 10 | -996252 | 2582.637 | -1253.09 | 321.75 | 0.124582 |
| 9 | 10 | -997183 | 1116.865 | -931.34 | 640.41 | 0.5734 |
| 10 | 10 | -997474 | 2154.987 | -290.93 | NA | NA |

Table S7. Results of Mantel tests for isolation by distance (IBD) using Nei’s genetic distance (D) for landscape samples and pairwise genetic differentiation (G_st_) for population samples. Landscape samples were partitioned between the zones of core, early expansion, late expansion, and invasion front to determine whether range expansion resulted in changes in isolation by distance. Correlation coefficient between the geographic distance and genetic distance for each of the IBD tests are included as R squared values.

| IBD Test | Observation | Std. Observation | Expectation | Variance | Simularted p-value | Rsquared |
| --- | --- | --- | --- | --- | --- | --- |
| Population Samples | 2.26E-02 | 1.59E-01 | 2.10E-04 | 1.66E-02 | 0.418 | 0.023 |
| Landscape Samples: Core Only | 1.79E-02 | 2.18E-01 | 6.31E-04 | 6.27E-03 | 0.418 | 0.018 |
| Landscape Samples: Core + Early Expansion | -7.72E-03 | -1.62E-01 | 9.43E-05 | 2.33E-03 | 0.570 | -0.010 |
| Landscape Samples: Core + Early Expansion + Late Expansion | -3.58E-02 | -7.60E-01 | 2.30E-04 | 2.25E-03 | 0.773 | -0.040 |
| Landscape Samples: Core + Early Expansion + Late Expansion + Invasion Front | -1.93E-02 | -4.39E-01 | 1.20E-03 | 2.17E-03 | 0.676 | 0.045 |

Table S8. Niche differentiation permutation tests for niche similarity, expansion, stability, and unfilling compared across the core, early expansion, late expansion, and invasion front.

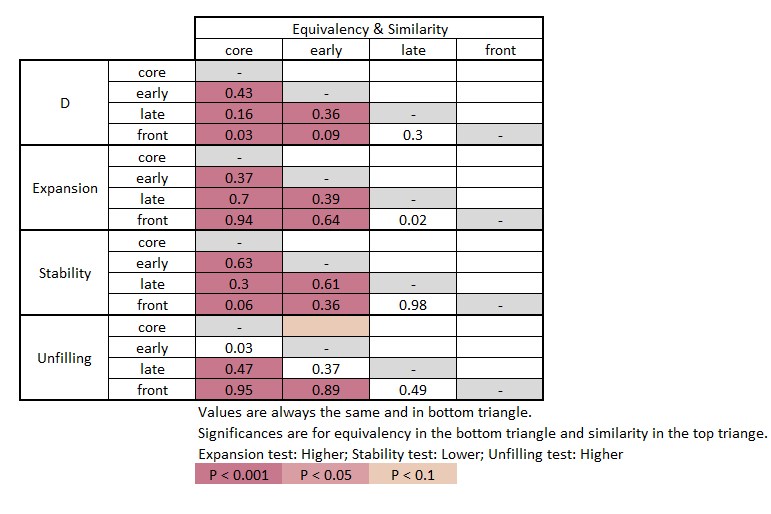

Table S9. Analysis of variance table for mixed effect models of germination probability. Significant terms are bolded.

| Mixed Model Anova Table (Type 3 tests, KR-method) | | | |
| --- | --- | --- | --- |
| Effect | F | Df | P |
| Temperature Regime | 26.29 | 4, 13.71 | **<0.001** |
| Source Temperature (above or below 20C) | 0.33 | 1, 12 | 0.579 |
| Temperature Regime : Source Temperature | 4.12 | 4, 392.57 | **0.003** |

| Mixed Model Anova Table (Type 3 tests, KR-method) | | | |
| --- | --- | --- | --- |
| Effect | F | Df | P |
| Temperature | 30.55 | 4, 11.11 | **<0.001** |
| Region | 0.07 | 1, 13 | 0.799 |
| Temperature:Region | 3.20 | 4, 423.67 | **0.013** |

Table S10. Tests for general linear contrasts between temperature treatment regimens and geographic regions using the ‘contrast.glht’ function from the R package ‘multcomp’.

| Simultaneous Tests for General Linear Hypotheses | | | | |
| --- | --- | --- | --- | --- |
| Linear Hypotheses: | Estimate | Std. Error | z value | Pr(>\|z\|) |
| Temp_5/15C == 0 | 0.55 | 0.64 | 0.86 | 0.771 |
| Temp_10/20C == 0 | 1.39 | 0.64 | 2.18 | 0.089 |
| Temp_15/25C == 0 | 0.05 | 0.64 | 0.08 | 1.000 |
| Temp_20/30C == 0 | 0.06 | 0.64 | 0.09 | 1.000 |
| Temp_25/35C == 0 | -0.38 | 0.64 | -0.59 | 0.928 |

Figure S1. Point map of leafy spurge ca. 1933. Hanson and Rudd (1933) documented in detail the distribution of leafy spurge across Minnesota and neighboring regions, providing a baseline for understanding the timeline of subsequent range expansion.

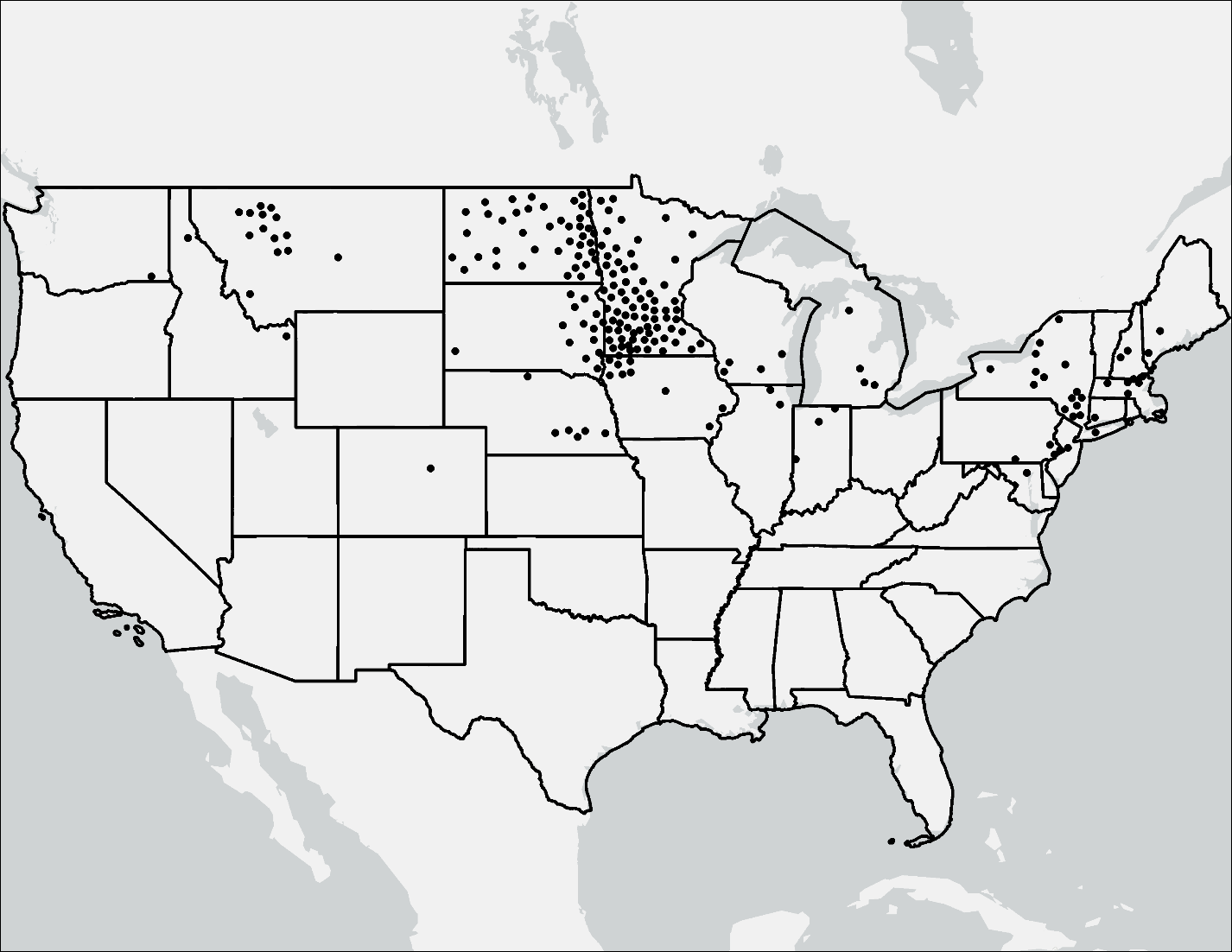

Figure S2. Filtering by minor allele frequency to remove homoeologous alleles. Leafy spurge is an auto-allohexaploid species so we expect likely homoeologous loci to have a 2:1 allelic ratio (e.g. AAAABB genotype). An excess of loci with an allele frequency around 0.33 was evident in our minor allele frequency spectrum. We removed loci with a minor allele frequency above 0.26 from our dataset because they are likely to have an excess of homoeologous genotypes.

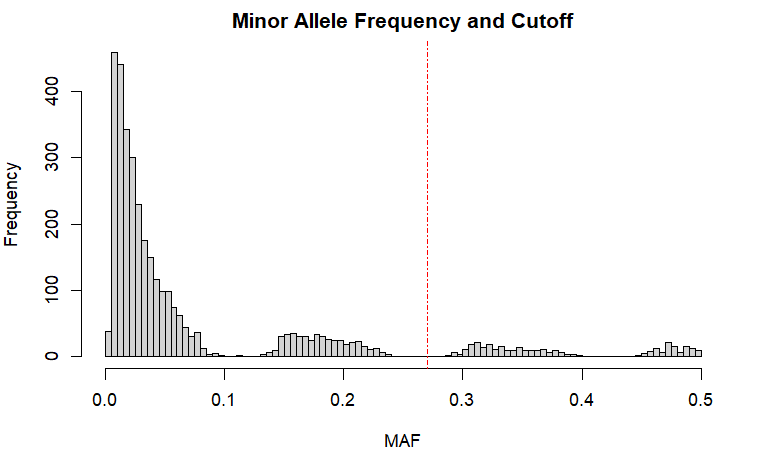

Figure S3. Residuals versus fitted plot for private alleles among populations.

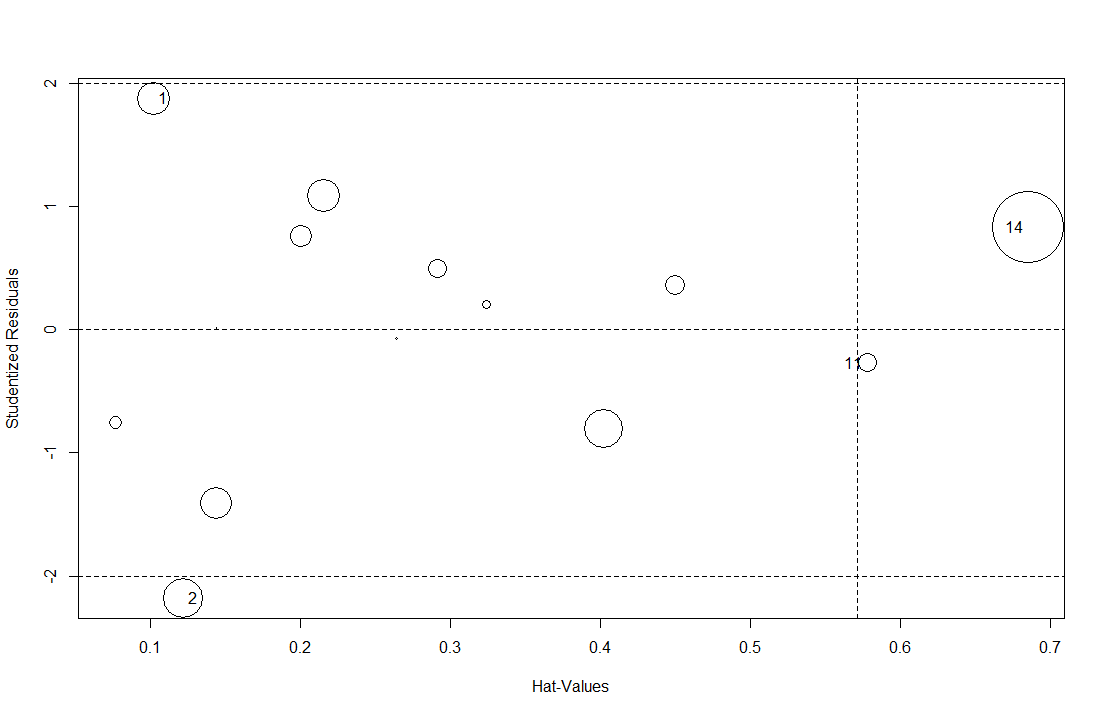

Plot of studentized residuals versus hat values for each of the population samples and the number of private alleles using the ‘influencePlot’ function from the R package ‘car’. The area of the circles represent observations proportional to the value Cook’s distance. Vertical line drawn at twice the average hat value, and horizontal reference lines at -2, 0, and 2 on the studentized-residual scale. Population number 14 is the Winona population, which was indicated as an outlier in the number of private alleles and was removed from the analysis of private alleles on latitude, longitude, and the interaction between the two variables.

Figure S4. PCA biplot of 19 BIOCLIM variables drawn from the landscape samples used in the ecological niche model (ENM). The PCA plot shows that there are three sets of highly correlated variables: variables on the bottom-right (Bio 1, 5, 6, 8, 10, 11) are largely related to temperature. Variables on the top-right are largely related to precipitation (Bio 12, 14, 16, 17, 18, 19). Variables on the left are largely related to ranges or CV (Bio 2, 3, 4, 7, and 15). The first two PCA axes explain 74.71% of variance.

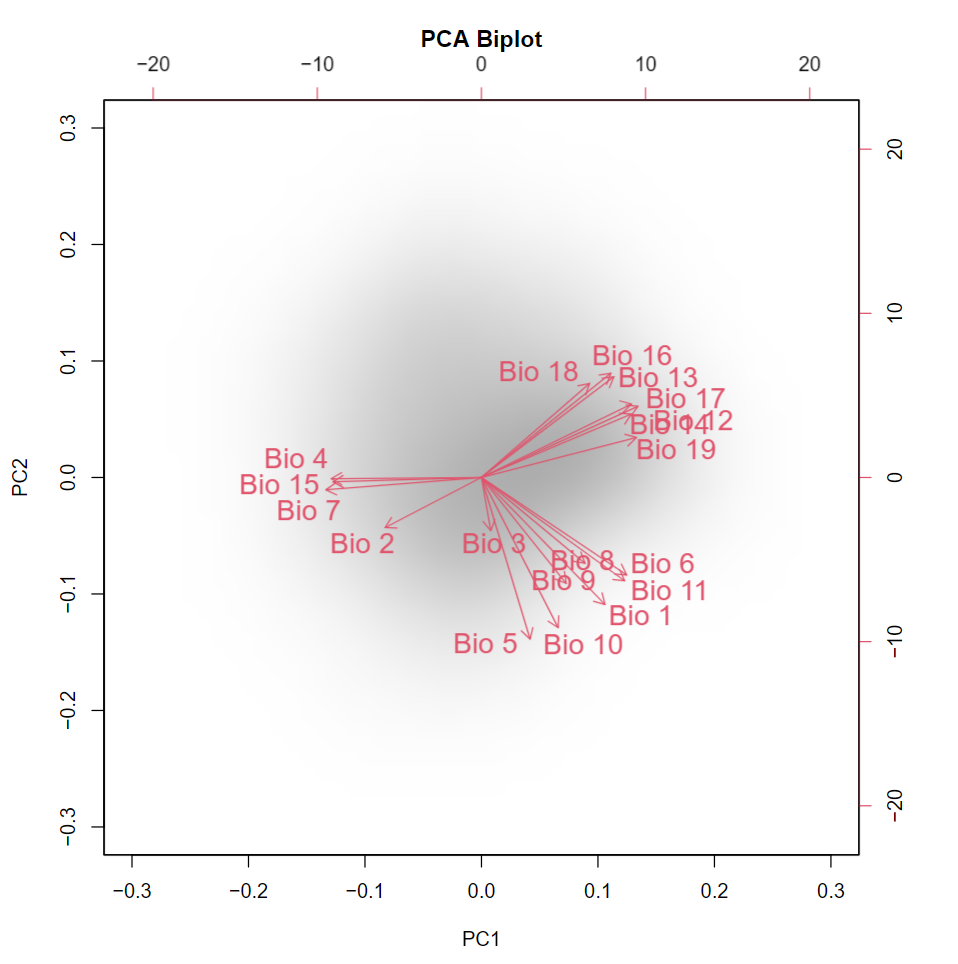

Figure S5. Contribution of environmental variables used in analyses of niche differentiation and ecological niche modeling as eigen vectors.

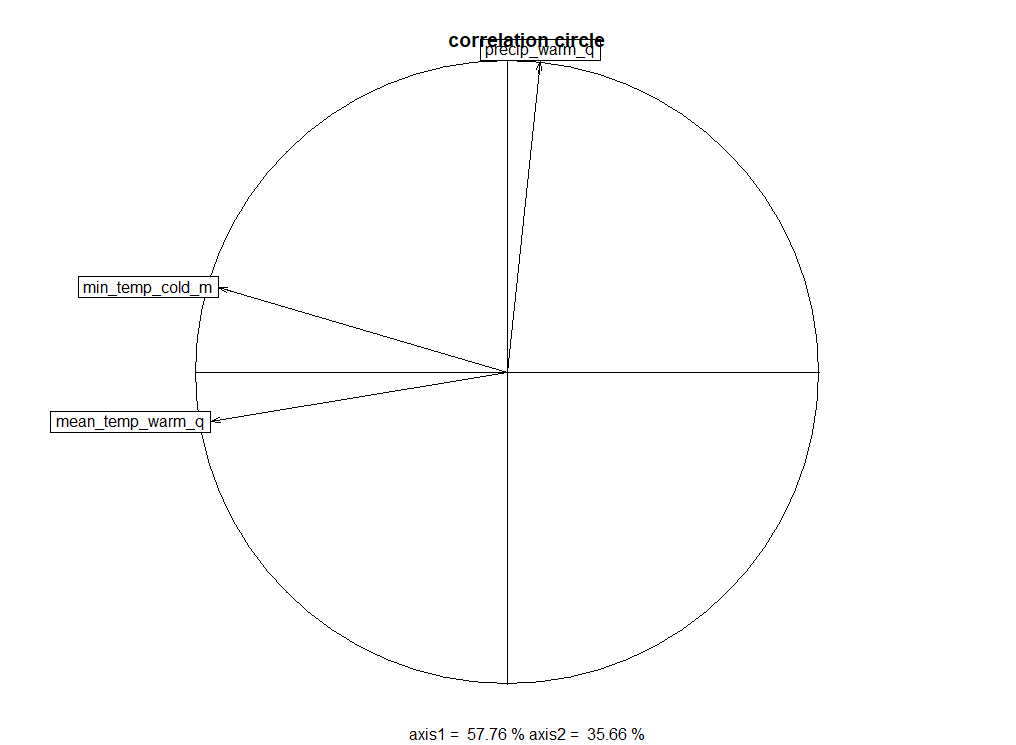

Figure S6. Environmental axes of niche differentiation between range core and invasion front.

Niche differences between the core and invasion front were most apparent along environmental axes related to temperature of the warmest quarter and minimum temperature of the coldest month, rather than precipitation. Green lines indicate environmental space sampled from the range core. Red lines indicate niche environmental sampled from the invasion front. Purple represents all available environmental space used in niche analyses.

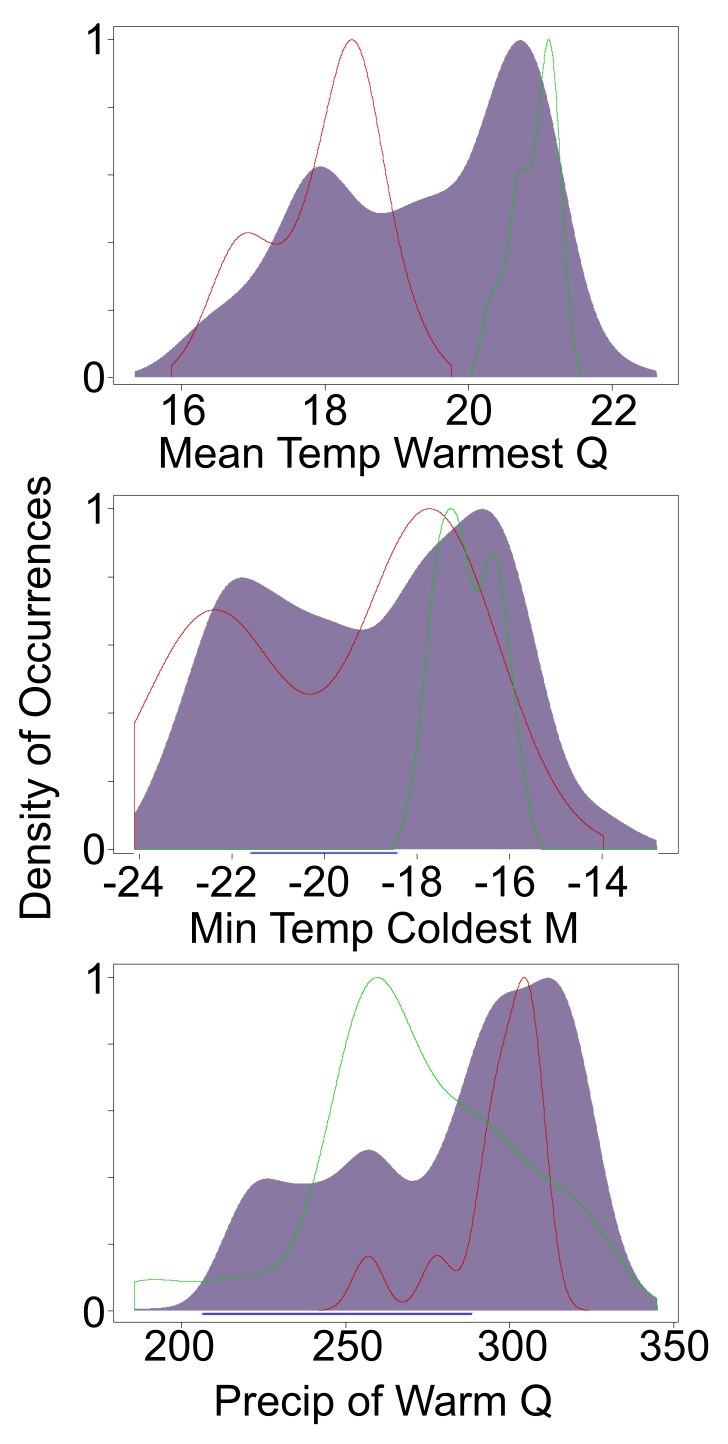

Figure S7. Variable response curves from the ecological niche model demonstrating variation in predicted habitat suitability (Y axis) given an environmental variable (X axis). Black lines indicate the model response to individual environmental variables. Yellow vertical lines indicate the position of seed sources from the germination experiment originating from areas with a mean temperature of the warmest quarter below 20C. Purple vertical lines indicate seeds sources originating from areas with a mean temperature of the warmest quarter above 20C. Red vertical line at 20C shows the bimodal separation of seed source populations by the temperature of the warmest quarter.

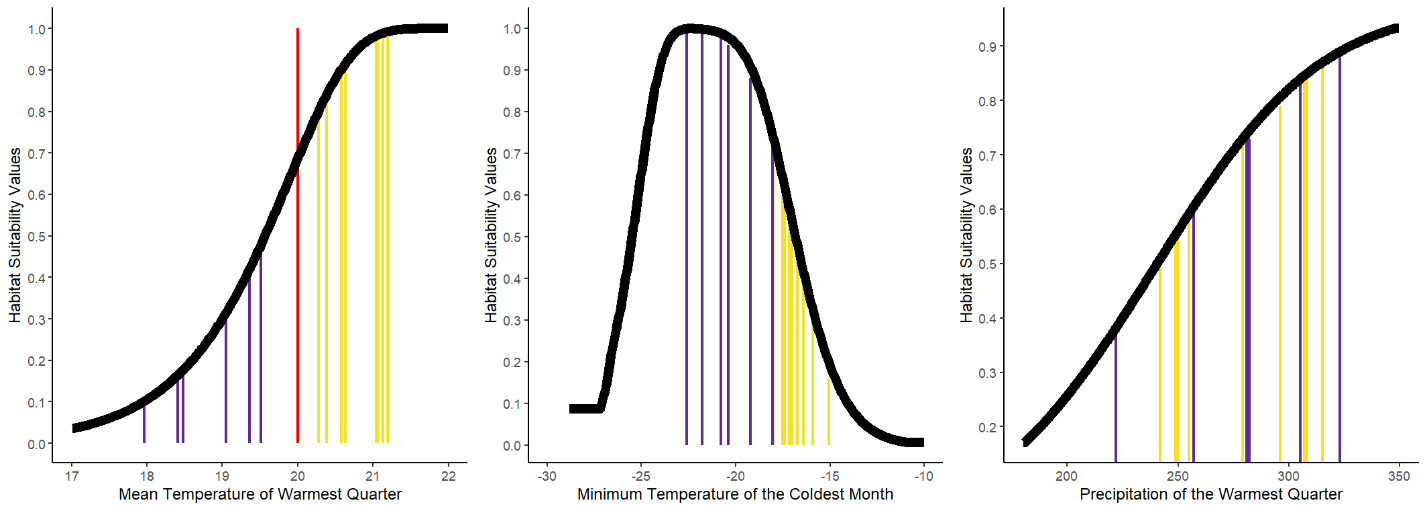

Figure S8. Scatterplots of predicted habitat suitability (Y axis) for each environmental variable (X axis) used in the ecological niche model. Blue line indicates linear model fit. Grey bars indicate confidence interval of fit. Points indicate occurrence samples used to fit the ecological niche model. Scatterplots demonstrate the correlation between predicted habitat suitability values and environmental variables.

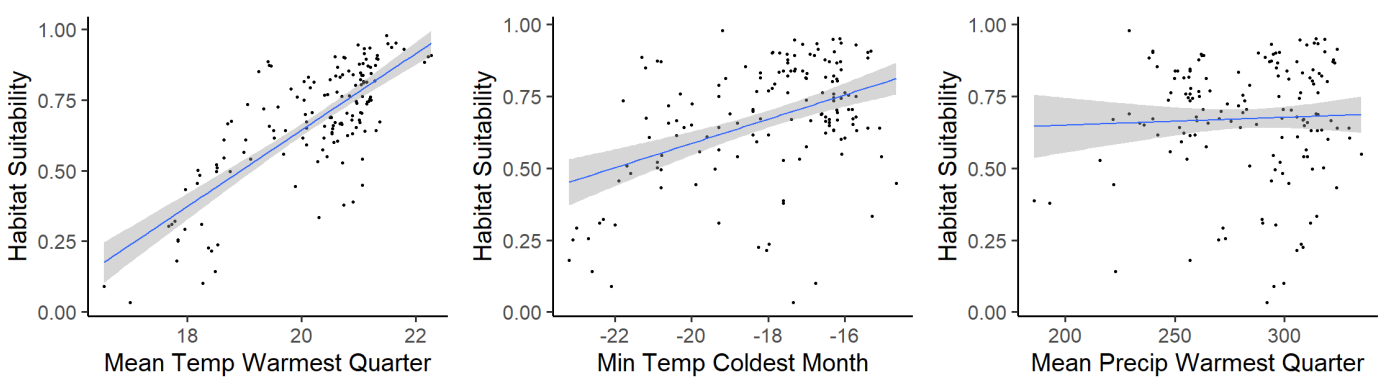
